# Pituitary stem cells produce paracrine WNT signals to control the expansion of their descendant progenitor cells

**DOI:** 10.1101/2020.05.22.107045

**Authors:** John Parkin Russell, Xinhong Lim, Alice Santambrogio, Val Yianni, Yasmine Kemkem, Bruce Wang, Matthew Fish, Scott Haston, Anaëlle Grabek, Shirleen Hallang, Emily Jane Lodge, Amanda Louise Patist, Andreas Schedl, Patrice Mollard, Roeland Nusse, Cynthia Lilian Andoniadou

## Abstract

In response to physiological demand, the pituitary gland generates new hormone-secreting cells from committed progenitor cells throughout life. It remains unclear to what extent pituitary stem cells (PSCs), which uniquely express SOX2, contribute to pituitary growth and renewal. Moreover, neither the signals that drive proliferation nor their sources have been elucidated. We have used genetic approaches in the mouse, showing that the WNT pathway is essential for proliferation of all lineages in the gland. We reveal that SOX2^+^ stem cells are a key source of WNT ligands. By blocking secretion of WNTs from SOX2^+^ PSCs *in vivo*, we demonstrate that proliferation of neighbouring committed progenitor cells declines, demonstrating that progenitor multiplication depends on the paracrine WNT secretion from SOX2^+^ PSCs. Our results indicate that stem cells can hold additional roles in tissue expansion and homeostasis, acting as paracrine signalling centres to coordinate the proliferation of neighbouring cells.

## INTRODUCTION

How stem cells interact with their surrounding tissue has been a topic of investigation since the concept of the stem cell niche was first proposed (Schofield, 1978). Secreted from supporting cells, factors such as WNTs, FGFs, SHH, EGF and cytokines, regulate the activity of stem cells (Nabhan et al., 2018; Palma et al., 2005; Tan and Barker, 2014). Furthermore, communication is known to take place in a bi-directional manner (Doupe et al., 2018; Tata and Rajagopal, 2016).

The anterior pituitary (AP) is a major primary endocrine organ that controls key physiological functions including growth, metabolism, reproduction and the stress responses and exhibits tremendous capability to remodel its constituent hormone populations throughout life, in response to physiological demand. It contains a population of *Sox2* expressing stem cells that self-renew and give rise to lineage-committed progenitors and functional endocrine cells (Andoniadou et al., 2013; Rizzoti et al., 2013). During embryonic development, SOX2^+^ undifferentiated precursor cells of Rathke’s pouch, the pituitary anlage (Arnold et al., 2011; Castinetti et al., 2011; Fauquier et al., 2008; Pevny and Rao, 2003), generate all committed endocrine progenitor lineages, defined by the absence of SOX2 and expression of either POU1F1 (PIT1), TBX19 (TPIT) or NR5A1 (SF1) (Bilodeau et al., 2009; Davis et al., 2011). These committed progenitors are proliferative and give rise to the hormone-secreting cells. Demand for hormone secretion rises after birth, resulting in dramatic organ growth and expansion of all populations by the second postnatal week (Carbajo-Perez and Watanabe, 1990; Taniguchi et al., 2002). SOX2^+^ pituitary stem cells (PSCs) are most active during this period, but the bulk of proliferation and organ expansion during postnatal stages derives from SOX2^-^ committed progenitors. The activity of SOX2^+^ PSCs gradually decreases and during adulthood is minimally activated even following physiological challenge (Andoniadou et al., 2013; Gaston-Massuet et al., 2011; Gremeaux et al., 2012; Zhu et al., 2015). By adulthood, progenitors carry out most of the homeostatic functions, yet SOX2^+^ PSCs persist throughout life in both mice and humans (Gonzalez-Meljem et al., 2017; Xekouki et al., 2018). The signals driving proliferation of committed progenitor cells are not known, and neither is it known if SOX2^*+*^ PSCs can influence this process beyond their minor contribution of new cells.

The self-renewal and proliferation of numerous stem cell populations relies upon WNT signals (Basham et al., 2019; Lim et al., 2013; Takase and Nusse, 2016; Wang et al., 2015; Yan et al., 2017). WNTs are necessary for the initial expansion of Rathke’s pouch as well as for PIT1 lineage specification (Osmundsen et al., 2017; Potok et al., 2008). In the postnatal pituitary, the expression of WNT pathway components is upregulated during periods of expansion and remodelling. Gene expression comparisons between neonatal and adult pituitaries or in GH-cell ablation experiments (Gremeaux et al., 2012; Willems et al., 2016), show that the WNT pathway is upregulated during growth and regeneration.

Our previous work revealed that during disease, the paradigm of supporting cells signalling to the stem cells may be reversed; mutant stem cells expressing a degradation-resistant β-catenin in the pituitary, promote cell non-autonomous development of tumours through their paracrine actions (Andoniadou et al., 2013; Gonzalez-Meljem et al., 2017). Similarly, degradation-resistant β-catenin expression in hair follicle stem cells led to cell non-autonomous WNT activation in neighbouring cells promoting new growth (Deschene et al., 2014). In the context of normal homeostasis, stem cells have been shown to influence daughter cell fate in the mammalian airway epithelium and the *Drosophila* gut via ‘forward regulation’ models, where the fate of a daughter cell is directed by a stem cell via juxtacrine Notch signalling (Ohlstein and Spradling, 2007; Pardo-Saganta et al., 2015). It remains unknown if paracrine stem cell action can also promote local proliferation in normal tissues.

Here, we used genetic approaches to determine if paracrine stem cell action takes place in the anterior pituitary and to discern the function of WNTs in pituitary growth. Our results demonstrate that postnatal pituitary expansion, largely driven by committed progenitor cells, depends on WNT activation. Importantly, we show that SOX2^+^ PSCs are the key regulators of this process, acting through secretion of WNT ligands acting in a paracrine manner on neighbouring progenitors. Identification of this forward-regulatory model elucidates a previously unidentified function for stem cells during tissue expansion.

## RESULTS

### WNT-responsive cells in the pituitary include progenitors driving major postnatal expansion

To clarify which cells respond to WNT signals in the postnatal anterior pituitary, we first characterised the anterior pituitary cell types activating the WNT pathway at P14, a peak time for organ expansion and a time point when a subpopulation of SOX2^+^ stem cells are proliferative. The *Axin2-CreERT2* mouse line (van Amerongen et al., 2012) has been shown to efficiently label cells with activated WNT signalling in the liver, lung, breast, skin, testes and endometrium among other tissues (Lim et al., 2013; Moiseenko et al., 2017; Syed et al., 2020; van Amerongen et al., 2012; Wang et al., 2015). *Axin2* positive cells were labelled by GFP following tamoxifen induction in *Axin2*^*CreERT2/+*^;*R26*^*mTmG/+*^ mice and pituitaries were analysed 2 days post-induction. We carried out double immunofluorescence staining using antibodies against uncommitted (SOX2), lineage committed (PIT1, TPIT, SF1), and hormone-expressing endocrine cells (GH, PRL, TSH, ACTH or FSH/LH) together with antibodies against GFP labelling the WNT-activated cells. We detected WNT-responsive cells among all the different cell types of the anterior pituitary including SOX2^+^ PSCs, the three committed populations and all hormone-secreting cells (Figure 1A, SFigure 1A).

**Figure 1.**
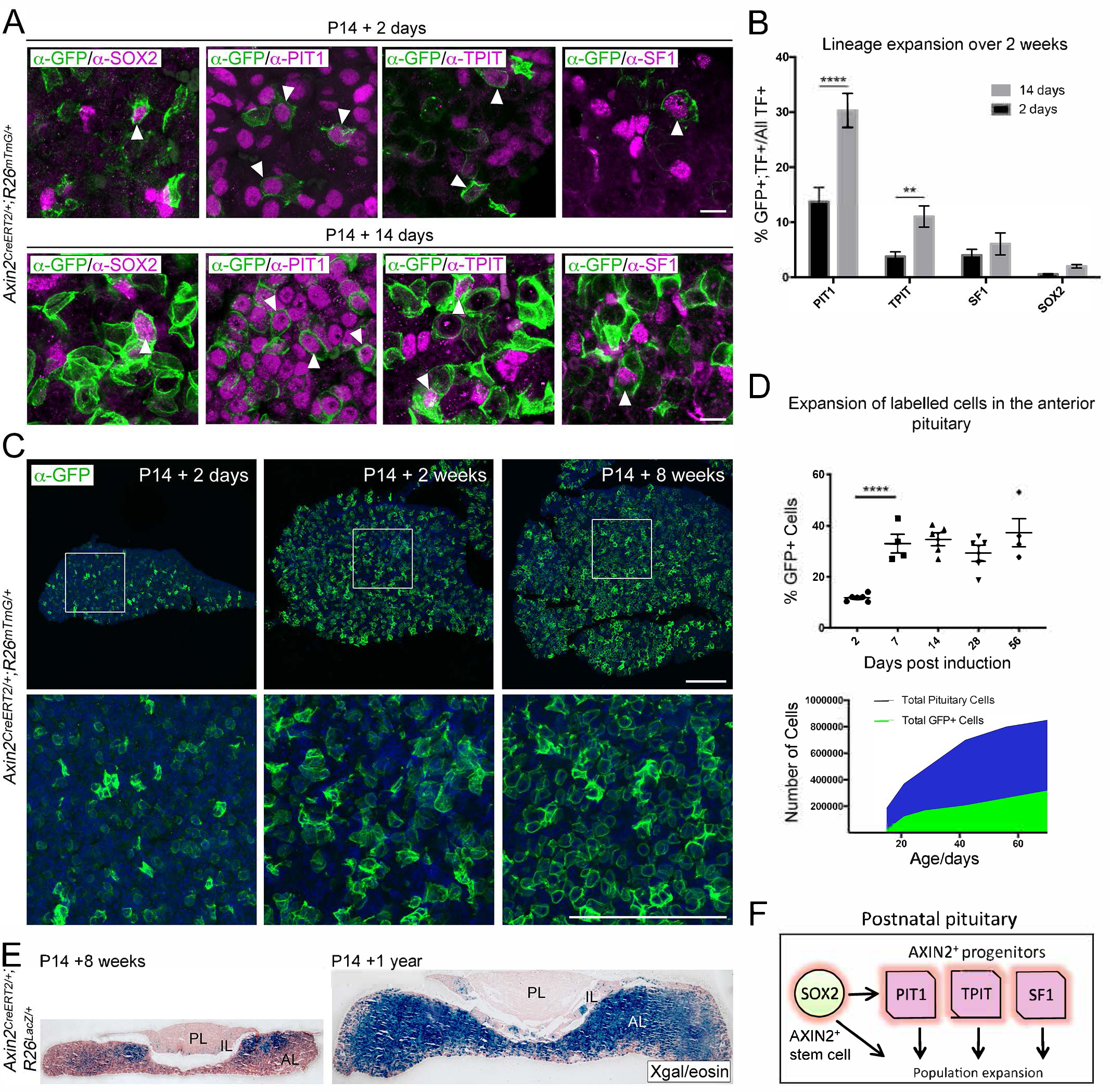
*Axin2* expressing cells contribute to pituitary growth and expansion of all lineages. A. Immunofluorescence staining against GFP (green) with markers of PSCs or lineage commitment (magenta) in *Axin2*^*CreERT2/+*^; *Rosa26*^*mTmG/+*^ pituitaries harvested from mice induced at P14 and lineage traced for 2 days (top panel) and 14 days (bottom panel). Scale bar 10αm. B. Quantification of lineage expansion between 2 and 14 days following induction at P14. Graph shows that the proportion of lineage committed cells (either PIT1^+^, TPIT^+^ or SF1^+^) and PSCs (SOX2^+^), i.e. that are transcription factor (TF)^+^ cells that are GFP^+^ increases between 2 days (black bars) and 14 days (grey bars) post induction. PIT1 *P*=0.000004, TPIT *P*=0.008 multiple *t*-tests. n = 4 animals per time point. C. Immunofluorescence staining against GFP (green) in pituitaries harvested from *Axin2*^*CreERT2/+*^;*Rosa26*^*mTmG/+*^ mice induced at P14 and lineage traced for 2 days, 2 weeks and 8 weeks. Bottom panel shows magnified fields of view of regions of interest indicated by white boxes in panels above. Scale bars 50αm. D. Top panel showing the quantification of the proportion of all cells in *Axin2*^*CreERT2/+*^;*Rosa26*^*mTmG/+*^ pituitaries that are GFP^+^ at 2, 7, 14, 28 and 56 days post induction as analysed by flow cytometry. Day 2 to 7 *P*<0.0001 unpaired *t*-test. (Bottom) Graph of the absolute number of GFP+ cells (green) and as a proportion of total cells (blue) at the time points indicated. E. X-gal staining in *Axin2*^*CreERT2/+*^;*Rosa26*^*LacZ/+*^ pituitaries harvested from mice induced at P14 and lineage traced for 8 weeks (left) and 1 year (right). F. Model summarising the major contribution of WNT-responsive progenitors of all lineages to pituitary growth, in addition to that of SOX2^+^ PSCs.

To confirm if the three committed lineages as well as uncommitted SOX2^+^ PSCs all expand in response to WNT, we further lineage-traced *Axin2*-expressing cells for 14 days after tamoxifen administration at P14. Double labelling revealed an increase in all four populations between 2 and 14 days (Figure 1A, B). This increase reached significance for the PIT1 (13.7% at 2 days to 30.3% at 14 days, *P*=0.000004) and TPIT (3.78% to 11.03%, *P*=0.008) populations, but not SF1 (0.5% to 4%, n.s.). Only a minority of SOX2^+^ PSCs were WNT-responsive at 2 days (0.57%) and this population expanded to 2% at 14 days (n.s.), suggesting that these are self-renewing. GFP^+^ cells were traced for a period of 8 weeks post-induction, which revealed that WNT-responsive descendants continued to expand at the same rate as the rest of the pituitary (n=4-8 mice per time point at P16, P21, P28, P42, P70) (Figure 1C, D). The time period between 2 and 7 days saw the greatest increase in GFP^+^ cells, during which, the labelled population nearly tripled in size (Figure 1D). The persistence of labelled cells was evident in longer-term traces using the *R26*^*lacZ/+*^ reporter (*Axin2*^*CreERT2/+*^;*R26*^*lacZ/+*^), up to a year following induction at P14 (Figure 1E, *n*=4). Clonal analysis using the Confetti reporter, demonstrated that individual *Axin2*-expressing cells (*Axin2*^*CreERT2/+*^;*R26*^*Confetti/+*^) gave a greater contribution after four weeks compared to lineage-tracing from *Sox2*-expressing cells (*Sox2*^*CreERT2/+*^;*R26*^*Confetti/+*^), in support of predominant expansion from WNT-responsive lineage-committed progenitors (SFigure 1B).

During times of greater physiological demand, the pituitary engages a proliferative response mirroring that observed during physiological growth (Levy, 2002; Nolan et al., 1998). To investigate if this is accompanied by an enhanced WNT response, we induced short-term physiological challenge through the induction of hypothyroidism in wild type juvenile mice for one week from P21. Feeding on an iodine-deficient diet supplemented with 0.15% propylthiouracil (PTU) for one week led to an increase in the number of new TSH-expressing thyrotrophs compared to control animals feeding on normal diet (SFigure 1C), as well as an increase in dividing cells marked by pH-H3 (SFigure 1D). This was accompanied by an elevation in *Axin2* mRNA transcripts (SFigure 1E) confirming an activation of the WNT response.

To establish if signalling mediated by β-catenin is necessary for organ expansion we carried out deletion of *Ctnnb1* in the *Axin2*^+^ population from P14 during normal growth (*Axin2*^*CreERT2/+*^;*Ctnnb1*^*lox(ex2-6)/lox(ex2-6)*^ hereby *Axin2*^*CreERT2/+*^;*Ctnnb1*^*LOF/LOF*^). Due to morbidity, likely due to *Axin2* expression in other organs, we were limited to analysis up to 5 days post-induction. This resulted in a significant reduction in the number of dividing cells lacking β-catenin, marked by pH-H3 (40% reduction, SFigure 1F, *P*=0.0313), confirming that activation of the WNT pathway is necessary for expansion of the pituitary populations. Taken together, these results confirm that postnatal AP expansion depends on WNT-responsive progenitors across all lineages, in addition to SOX2^+^ PSCs (Figure 1F).

### WNT/β-catenin signalling is required for long-term anterior pituitary expansion from SOX2^+^ pituitary stem cells

We further explored the role of WNT pathway activation in postnatal SOX2^+^ stem cells. To permanently mark WNT-responsive cells and their descendants whilst simultaneously marking SOX2^+^ PSCs, we combined the tamoxifen-inducible *Axin2*^*CreERT2/+*^;*R26*^*tdTomato/+*^ with the *Sox2-Egfp* strain, where cells expressing SOX2 are labelled by EGFP (*Axin2*^*CreERT2/+*^;*Sox2*^*Egfp/+*^;*R26*^*tdTomato/+*^). Following tamoxifen administration from P21, tdTomato- and EGFP-labelled cells were analysed by flow sorting after 72h, by which point all induced cells robustly express tdTomato (Figure 2A). Double-labelled cells comprised 1.5-2% of the SOX2^+^ population (Figure 2A, arrowheads), with the majority of tdTomato^+^ cells found outside of the SOX2^+^ compartment. It was previously shown that only around 2.5-5% of SOX2^+^ PSCs have clonogenic potential through *in vitro* assays (Andoniadou et al., 2012; Andoniadou et al., 2013; Perez Millan et al., 2016). To determine if WNT-responsive SOX2^+^ cells are stem cells capable of forming colonies, we isolated double positive tdTomato^+^;EGFP^+^ cells (i.e. *Axin2*^*+*^;*Sox2*^*+*^) as well as the single-expressing populations and plated these in equal numbers in stem cell-promoting media at clonal densities (Figure 2B). Double positive tdTomato^+^;EGFP^+^ cells showed a significant increase in the efficiency of colony formation compared to single-labelled EGFP^+^ cells (average 9% compared to 5%, *n*=5 pituitaries, *P*=0.0226, Mann-Whitney *U* test (two-tailed)), demonstrating WNT-responsive SOX2^+^ PSCs have a greater clonogenic potential under these *in vitro* conditions, confirming *in vivo* data in Figure 1B. As expected from previous work, none of the single-labelled tdTomato^+^ cells (i.e. SOX2 negative) were able to form colonies (Andoniadou et al., 2012).

**Figure 2.**
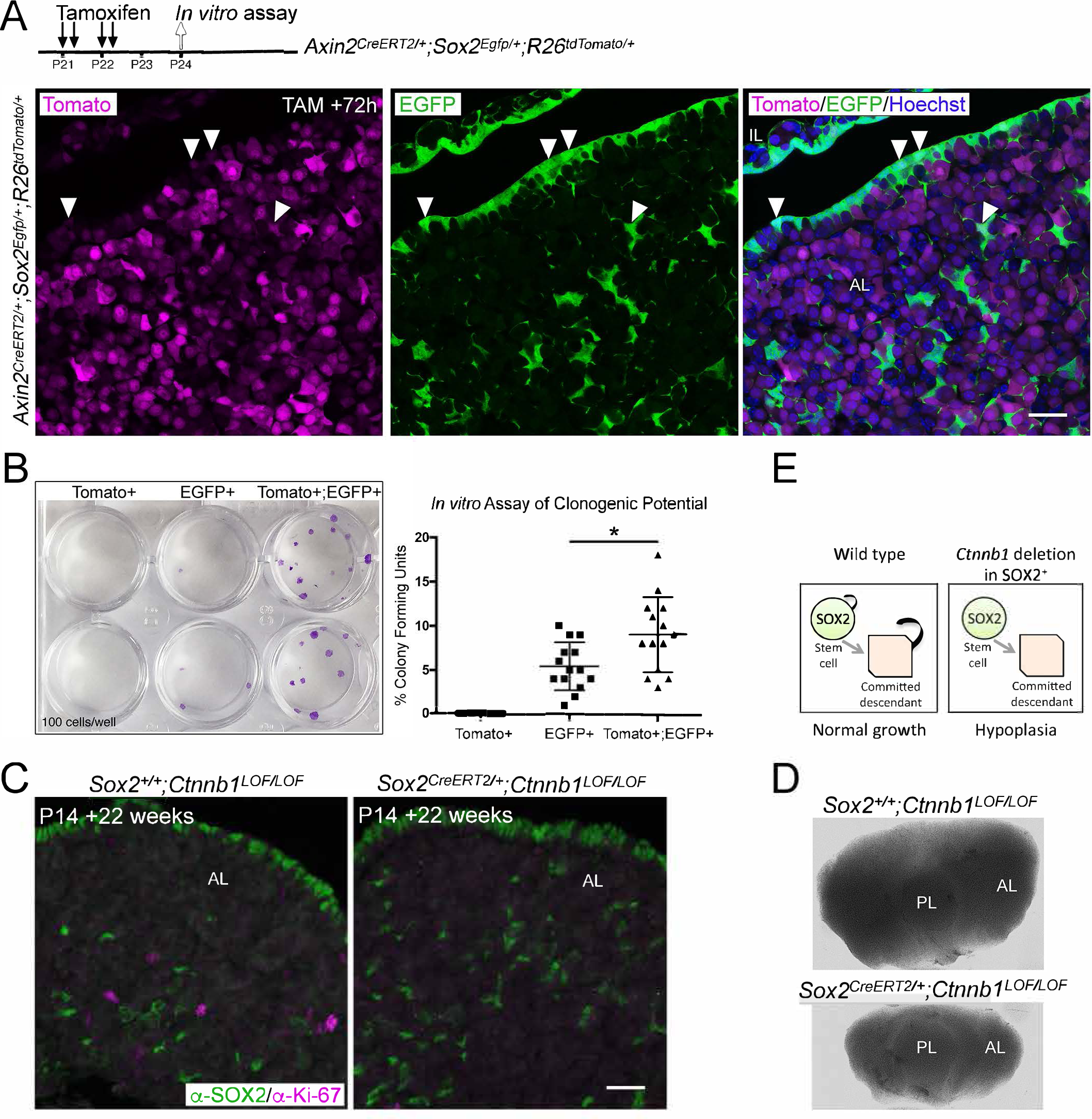
Activation of WNT signalling in SOX2^+^ PSCs and their descendants is necessary for long-term growth. A. Schematic of the experimental timeline used in panels A and B. Endogenous expression of tdTomato (magenta, *Axin2* targeted cells) and EGFP (green, *Sox2* expressing cells) in *Axin2*^*CreERT2/+*^;*Sox2*^*Egfp/+*^;*Rosa26*^*tdTomato/+*^ pituitaries harvested at P24 sectioned in the frontal plane. Nuclei are counterstained with Hoechst in the merged panel. Scale bar 50αm B. A representative culture plate showing colonies derived from Tomato^+^, EGFP^+^ or Tomato^+^;EGFP^+^ cells that were isolated from *Axin2*^*CreERT2/+*^;*Sox2*^*Egfp/+*^;*Rosa26*^*tdTomato/+*^ pituitaries by FACS plated in stem cell promoting media at clonogenic densities and stained with crystal violet (left panel). The proportion of colony-forming cells in each subpopulation were quantified by counting the number of colonies per well (right panel). Each data point indicates individual wells. *P*=0.0226, Mann-Whitney *U* test (two-tailed). C. Immunofluorescence staining against SOX2 (green) and Ki-67 (magenta) in *Sox2*^*+/+*^*Ctnnb1*^*LOF/LOF*^ (control) and *Sox2*^*CreERT2/+*^*Ctnnb1*^*LOF/LOF*^ (mutant) pituitaries from mice induced at P14 and analysed 22 weeks after induction (at P168) (bottom panel). Scale bar 50αm. D. Dorsal view of whole mount pituitaries of *Sox2*^*+/+*^;*Ctnnb1*^*LOF/LOF*^ (control) and *Sox2*^*CreERT2/+*^;*Ctnnb1*^*LOF/LOF*^ (mutant), 22 weeks after induction (i.e. P168). E. Model summarising the effect of *Ctnnb1* deletion in SOX2^+^ PSCs. PL, posterior lobe; IL, intermediate lobe; AL, anterior lobe. All plotted points equal one technical replicate, n = 5 biological replicates.

To confirm that PSCs with active WNT signalling through β-catenin have a greater propensity to form colonies *in vitro*, we analysed postnatal pituitaries from *TCF/Lef:H2B-EGFP* mice, reporting the activation of response to WNT signals. This response is detected through expression of an EGFP-tagged variant of histone H2B, which is incorporated into chromatin (Ferrer-Vaquer et al., 2010). At P21, EGFP^+^ cells were abundant in all three lobes and particularly in the marginal zone harbouring SOX2^+^ stem cells (SFigure 2A). Through double mRNA *in situ* hybridisation against *Egfp* and *Sox2* in *TCF/Lef:H2B-EGFP* pituitaries, we confirmed that *Sox2*-expressing cells activate H2B-EGFP expression at this time point (SFigure 2B). Isolation by fluorescence-activated cell sorting and *in vitro* culture of the postnatal EGFP^+^ compartment revealed an enrichment of cells with clonogenic potential in the EGFP^High^ fraction compared to EGFP^Low^ or negative cells (SFigure 2C, *n*=5 pituitaries). Together these results reveal that a proportion of postnatal SOX2^+^ stem cells respond to WNTs through downstream β-catenin/TCF/LEF signalling and that these cells have greater clonogenic capacity *in vitro*.

To further address the role of the canonical WNT response in the activity of SOX2^+^ PSCs *in vivo*, we expressed a loss-of-function allele of β-catenin specifically in *Sox2*-expressing cells (*Sox2*^*CreERT2/+*^;*Ctnnb1*^*lox(ex2-6)/lox(ex2-6)*^ hereby *Sox2*^*CreERT2/+*^;*Ctnnb1*^*LOF/LOF*^) from P14. Twenty-two weeks following induction at P168, there was a substantial drop in the number of cycling cells in the pituitary of *Sox2*^*CreERT2/+*^;*Ctnnb1*^*LOF/LOF*^ mutants compared to *Sox2*^*+/+*^;*Ctnnb1*^*LOF/LOF*^ controls (Figure 2C, *n*=2 pituitaries per genotype). This was accompanied by anterior pituitary hypoplasia following the loss of *Ctnnb1* in SOX2^+^ PSCs (Figure 2D). Therefore, the proliferative capacity of *Ctnnb1*-deficient SOX2^+^ PSCs and of their descendants was impaired long-term, leading to reduced growth. *In vivo* genetic tracing of targeted cells over the 22-week period (*Sox2*^*CreERT2/+*^;*Ctnnb1*^*LOF/+*^;*R26*^*mTmG/+*^ compared to *Sox2*^*CreERT2/+*^;*Ctnnb1*^*LOF/LOF*^;*R26*^*mTmG/+*^ pituitaries) revealed that targeted (*Ctnnb1*-deficient) SOX2^+^ PSCs were capable of giving rise to the three committed lineages PIT1, TPIT and SF1 (SFigure 2D), indicating that the loss of *Ctnnb1* does not prevent differentiation of SOX2^+^ PSCs into the three lineages. Downregulation of β-catenin was confirmed by immunofluorescence in SOX2^+^ (mGFP^+^) derivatives (SFigure 2E). In conclusion, WNT/β-catenin signalling is cell-autonomously required to promote the expansion of all pituitary populations (Figure 2E).

### SOX2^+^ stem cells express WNT ligands

Having established that WNT activation is responsible for promoting proliferation in the AP, we next focused on identifying the source of WNT ligands. *Axin2* expressing cells from *Axin2*^*CreERT2/+*^;*R26*^*mTmG/+*^ mice were labelled at P14 by tamoxifen induction. Cells expressing *Axin2* at the time of induction are labelled by GFP expression in the membrane. Double immunofluorescence staining for GFP together with SOX2 revealed that *Axin2* expressing cells (mGFP^+^) are frequently located in close proximity to SOX2^+^ PSCs (Figure 3A). Two-dimensional quantification of the two cell types revealed that over 50% of mGFP^+^ cells were in direct contact with SOX2^+^ nuclei (*n*=3 pituitaries, >500 SOX2^+^ cells per gland, Figure 3A). The analysis did not take into account the cellular processes of SOX2^+^ cells. These results led us to speculate that SOX2^+^ PSCs may be a source of key WNT ligands promoting proliferation of lineage-committed cells.

**Figure 3.**
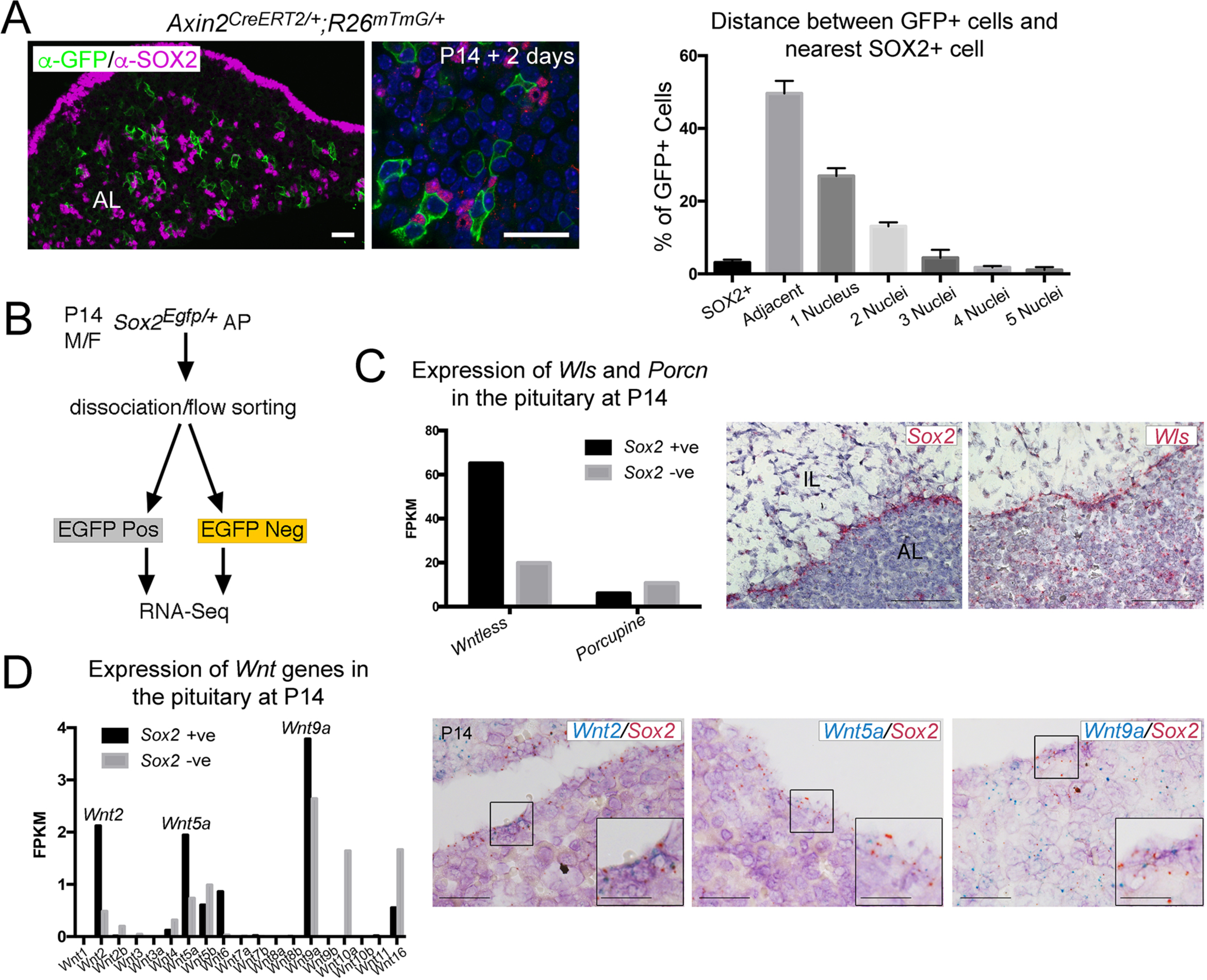
SOX2^+^ PSCs are as a source of WNT ligands in the pituitary. A. Immunofluorescence staining against GFP (green) and SOX2 (magenta) in *Axin2*^*CreERT2/+*^; *Rosa26*^*mTmG/+*^ pituitaries 48 hours post induction. Graph representing a quantification of the proximity of individual GFP^+^ cells to the nearest SOX2^+^ cell as quantified by the number of nuclei separating them. Scale bars 50αm. B. Experimental paradigm for RNA Seq analysis of *Sox2* positive and negative cells. C. Graphs representing the FPKM values of *Wls* and *Porcupine* in *Sox2* positive and negative cells (black and grey bars, respectively). mRNA in situ hybridisation for *Sox2* and for *Wls* on wild type sagittal pituitaries at P14, demonstrating strong *Wls* expression in the marginal zone epithelium. Scale bars 250αm. D. Bar chart showing the FPKM values of *Wnt* genes in the *Sox2+* and *Sox2-* fractions. Double mRNA in situ hybridisation against *Wnt2, Wnt5a* and *Wnt9a* (blue) together with *Sox2* (red) validating expression in the *Sox2+* population. Boxed regions through the marginal zone epithelium are magnified. Scale bars 100αm and 50αm in boxed inserts.

In order to determine if SOX2^+^ PSCs express WNT ligands, we carried out gene expression profiling of SOX2^+^ and SOX2^-^ populations at P14, through bulk RNA-sequencing. Pure populations of *Sox2*-expressing cells excluding lineage-committed populations, were isolated from *Sox2*^*Egfp/+*^ male and female pituitaries at P14 based on EGFP expression as previously shown (Andoniadou et al., 2012) (Figure 3B, SFigure 3A). Analysis of global gene expression signatures using ‘Gene Set Enrichment Analysis’ (GSEA) (Subramanian et al., 2005) identified a significant enrichment of molecular signatures related to EMT, adherens and tight junctions in the EGFP^+^ fraction, characteristic of the SOX2^+^ population (SFigure 3B). The SOX2^+^ fraction also displayed enrichment for genes associated with several signalling pathways known to be active in these cells, including EGFR (Iwai-Liao et al., 2000), Hippo (Lodge et al., 2016; Lodge et al., 2019; Xekouki et al., 2019), MAPK (Haston et al., 2017), FGF (Higuchi et al., 2017), Ephrin (Yoshida et al., 2015; Yoshida et al., 2017) and p53 (Gonzalez-Meljem et al., 2017) (SFigure 3C, Supplementary Table 1). Additionally, PI3K, TGFβ and BMP pathway genes were significantly enriched in the SOX2^+^ population (SFigure 3C, Supplementary Table 1). Query of the WNT-associated genes did not suggest a global enrichment in WNT targets (e.g. enrichment of *Myc* and *Jun*, but not of *Axin2* or *Lef1*) (SFigure 3D, Supplementary Table 1). Instead, SOX2^+^ PSCs expressed a unique transcriptomic fingerprint of key pathway genes including *Lgr4, Znrf3, Rnf43* capable of regulating WNT signal intensity in SOX2^+^ PSCs, as well as enriched expression of the receptors *Fzd1, Fzd3, Fzd4, Fzd6* and *Fzd7* (SFigure 3D). The predominant R-spondin gene expressed in the pituitary was *Rspo4*, specifically by the EGFP-negative fraction (SFigure 3D). The gene profiling revealed that *Wls* expression, encoding Gpr177/WLS, a necessary mediator of WNT ligand secretion (Carpenter et al., 2010; Takeo et al., 2013; Wang et al., 2015), is enriched in SOX2^+^ PSCs (Figure 3C). Analysis of *Wnt* gene expression confirmed enriched expression of *Wnt2, Wnt5a* and *Wnt9a* in SOX2^+^ PSCs, and the expression of multiple additional *Wnt* genes by both fractions at lower levels (SOX2^+^ fraction: *Wnt5b, Wnt6, Wnt16*; SOX2^-^ fraction: *Wnt2, Wnt2b, Wnt3, Wnt4, Wnt5a, Wnt5b, Wnt9a, Wnt10a, Wnt16*) (Figure 3D). These results reveal that SOX2^+^ PSCs express the essential components to regulate activation of the WNT pathway and express *Wnt* genes as well as the necessary molecular machinery to secrete WNT ligands.

### Paracrine signalling from SOX2^+^ stem cells promotes WNT activation

We sought to conclusively determine if WNT secretion specifically from SOX2^+^ PSCs drives proliferation of surrounding cells in the postnatal pituitary gland. We proceeded to delete *Wls* only in the *Sox2*-expressing population (*Sox2*^*CreERT2/+*^;*Wls*^*fl/fl*^) from P14 by a series of tamoxifen injections. Due to morbidity, we limited analyses to one week following induction. Pituitaries appeared mildly hypoplastic at P21 along the medio-lateral axis (SFigure 4). To determine if this is a result of reduced proliferation, we carried out immunofluorescence using antibodies against Ki-67 and SOX2. This revealed significantly fewer cycling cells in the SOX2^-^ population of *Sox2*^*CreERT2/+*^;*Wls*^*fl/fl*^ mutant pituitaries compared to *Sox2*^*+/+*^;*Wls*^*fl/fl*^ controls (10.326% Ki-67 in control compared to 3.129% in mutant, *P*=0.0008, unpaired *t*-test) (Figure 4A). Additionally, we observed a reduction of cycling cells within the SOX2^+^ population (5.582% Ki-67 in control compared to 2.225% in induced *Sox2*^*CreERT2/+*^;*Wls*^*fl/fl*^ mutant pituitaries, *P*=0.0121, unpaired *t*-test) (Figure 4A). To determine if reduced levels of WNT activation accompanied this phenotype, we carried out double mRNA *in situ* hybridisation using specific probes against *Lef1* and *Sox2*. There was an overall reduction in *Lef1* expression in mutants compared to controls, in which we frequently observed robust expression of *Lef1* transcripts in close proximity to cells expressing *Sox2* (arrows, Figure 4B). Together, our data support a paracrine role for SOX2^+^ pituitary stem cells in driving the expansion of committed progeny through the secretion of WNT ligands (Figure 4C).

**Figure 4.**
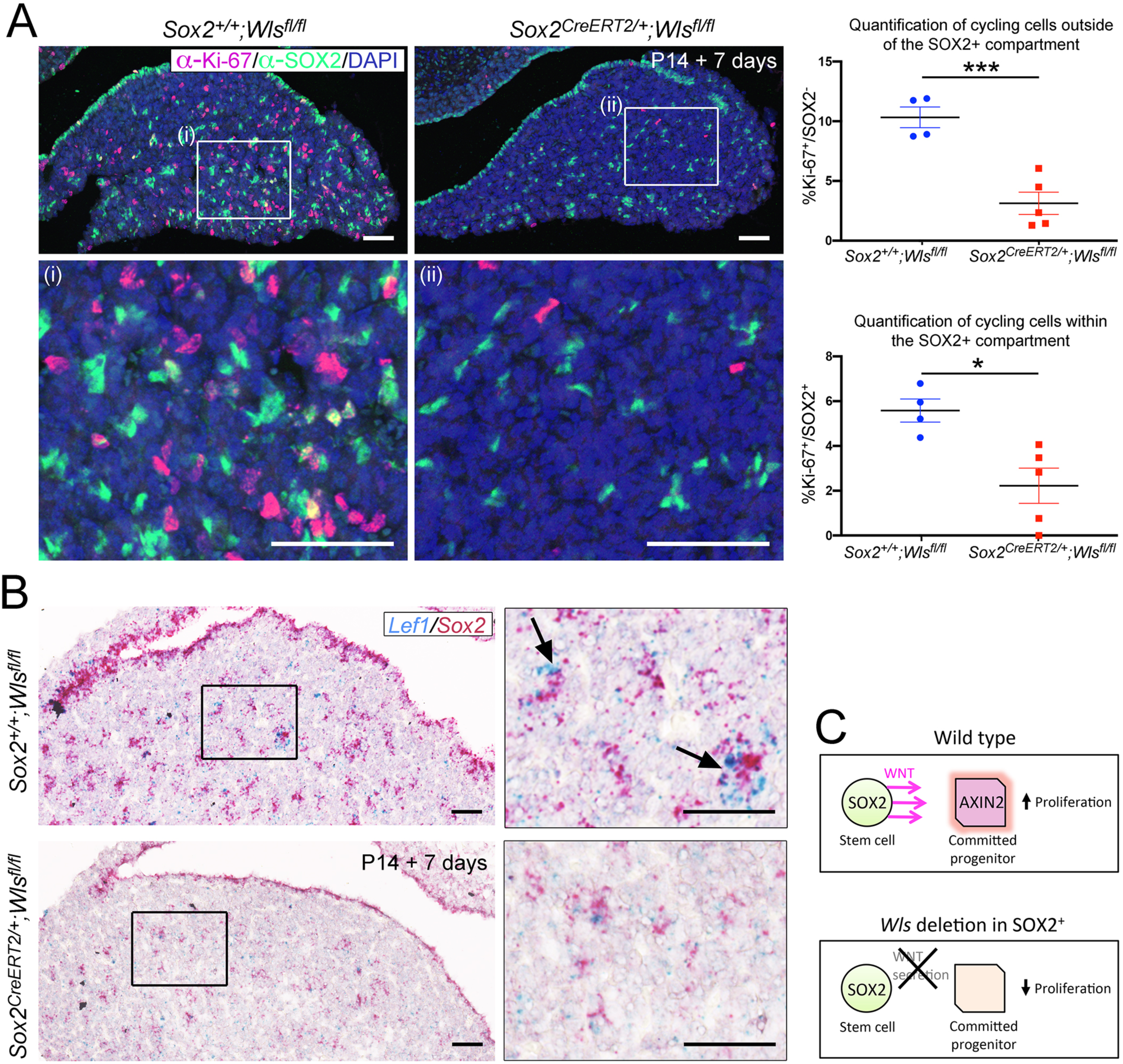
Paracrine secretion of WNTs from SOX2^+^ PSCs is necessary for expansion of committed cells. A. Immunofluorescence staining against SOX2 (green) and Ki-67 (magenta) in *Sox2*^*+/+*^;*Wls*^*fl/fl*^ (control) and *Sox2*^*CreERT2/+*^;*Wls*^*fl/fl*^ (mutant) pituitaries induced from P14 and analysed after one week. Nuclei were counterstained with Hoechst. (i) and (ii) represent magnified fields of view of regions indicated by white boxes in top panels. Scale bars 50αm. Graph of quantification of cycling cells marked by Ki-67 among cells negative for SOX2. Values represent mean +/-SEM, *P*=0.0008, unpaired *t*-test. Graph of quantification of cycling cells marked by Ki-67 among SOX2-positive cells. Values represent mean +/-SEM, *P*=0.0121, unpaired *t*-test. B. Double mRNA in situ hybridisation using specific probes against *Lef1* (blue) and *Sox2* (red) in control and mutant pituitaries following tamoxifen induction from P14 and tracing for 7 days. Scale bars 250αm and 50αm in boxed regions. C. Model summarising paracrine WNT secretion from SOX2^+^ PSCs to lineage-committed progenitors and the effects of abolishing WNT secretion from SOX2^+^ PSCs through the deletion of *Wls*.

## DISCUSSION

Emerging disparities between the archetypal stem cell model, exhibited by the haematopoietic system, and somatic stem cells of many organs, have led to the concept that stem cell function can be executed by multiple cells not fitting a typical stem cell paradigm (Clevers and Watt, 2018). In organs with persistent populations possessing typical functional stem cell properties yet contributing minimally to turnover and repair, the necessity for such classical stem cells is questioned. Here we show that WNT signalling is required for postnatal pituitary growth by both SOX2^+^ PSCs as well as SOX2^-^ committed progenitors. We identify an additional discreet function for SOX2^+^ PSCs, where these signal in a feedforward manner by secreting WNT ligands as cues to stimulate proliferation and promote tissue growth.

Consistent with previous reports, our data support that SOX2^+^ PSCs contribute, but do not carry out the majority of tissue expansion during the postnatal period (Zhu et al., 2015); instead, new cells primarily derive from more committed progenitors, which we show to be WNT-responsive. We demonstrate that this population of lineage-restricted WNT-responsive cells rapidly expands and contributes long-lasting clones from postnatal stages. It remains to be shown if cells among the SOX2^-^ lineage-committed populations may also fall under the classical definition of a stem cell. Preventing secretion of WNT ligands from SOX2^+^ PSCs reveals that far from being dispensable, paracrine actions of the SOX2^+^ population that are inactive in their majority, are necessary for anterior lobe expansion from lineage-committed populations. In the adrenal, R-spondins are necessary for cortical expansion and zonation, where deletion of *Rspo3*, expressed by the capsule which contains adrenocortical stem cells, results in reduced proliferation of the underlying steroidogenic cells (Vidal et al., 2016). Corroborating a model where committed pituitary progenitors depend on the paracrine actions of SOX2^+^ PSCs, Zhu and colleagues observed that in pituitaries with reduced numbers of PSCs, proliferation among PIT1^+^ cells was significantly impaired (Zhu et al., 2015). It would be intriguing to see if there is a reduction in WNT signalling in this model, or following genetic ablation of adult SOX2^+^ PSCs (Roose et al., 2017).

We show that a sub-population of SOX2^+^ PSCs in the postnatal gland are also WNT-responsive and have greater *in vitro* colony-forming potential under defined conditions. This colony-forming potential is normally a property of a minority of SOX2^+^ PSCs at any given age and reflects their *in vivo* proliferative capacity (Andoniadou et al., 2012; Rizzoti et al., 2013). A role for the WNT pathway in promoting SOX2^+^ cell activity is supported by studies showing that pathogenic overexpression of β-catenin promotes their colony-forming ability (Sarkar et al., 2016), and their *in vivo* expansion (Andoniadou et al., 2012). Additionally, elevated WNT pathway activation has been described for pituitary side-population cells, enriched for SOX2^+^ stem cells from young, compared to old pituitaries (Gremeaux et al., 2012). This is in line with our findings that the WNT pathway has an important function in promoting the activation of SOX2^+^ PSCs. It remains to be shown if this response relies on autocrine WNT-signalling as for other stem cells (Lim et al., 2013), however our results reveal reduced proliferation among SOX2^+^ PSCs when WNT secretion from these cells is abolished, supportive of either autocrine signalling, or paracrine signalling between different subsets of the SOX2^+^ population.

The mechanism preventing the majority of SOX2^+^ PSCs from responding to WNT signals remains elusive but points to heterogeneity among the population. Such regulation could occur at the level of receptor signalling; we have shown by bulk transcriptomic profiling that SOX2^+^ PSCs express the receptors required to respond to the WNT pathway, but also express high levels of the frizzled inhibitor *Znrf3*, and the R-spondin receptor *Lgr4*. One conceivable scenario is that high levels of *Znrf3* inhibit frizzled receptors in the absence of R-spondin under normal physiological conditions, supressing a WNT response. In support of this, R-spondins have been shown to promote pituitary organoid formation (Cox et al., 2019). Whether the R-spondin/LGR/ZNRF3 module is active under physiological conditions needs to be determined. Furthermore, well-described factors expressed in PSCs are known to have inhibitory effects on β-catenin-mediated transcription, such as YAP/TAZ (Azzolin et al., 2014; Gregorieff et al., 2015) and SOX2 itself (Alatzoglou et al., 2011; Kelberman et al., 2008).

In summary, we demonstrate an alternative mechanism for stem cell contribution to homeostasis, whereby these can act as paracrine signalling hubs to promote local proliferation. Applicable to other organs, this missing link between SOX2^+^ PSCs and committed cell populations of the anterior pituitary, is key for basic physiological functions and renders stem cells integral to organ expansion.

## MATERIALS AND METHODS

### Mice

All procedures were performed under compliance of the Animals (Scientific Procedures) Act 1986, Home Office License and KCL Ethical Review approval. KCL Biological Services Unit staff undertook daily animal husbandry. Genotyping was performed on ear biopsies taken between P11 and P15 by standard PCR using the indicated primers. These experiments were not conducted at random and the experimenters were not blind while conducting the animal handling and assessment of tissue. Images are representative of the respective genotypes.

For lineage tracing studies male *Axin2*^*CreERT2/+*^ or *Sox2*^*CreERT2/+*^ mice were bred with homozygous *ROSA26*^*mTmG/mTmG*^ or *ROSA26*^*Confetti/Confetti*^ dams to produce the appropriate allele combinations on the reporter background. Pups were induced at P14 or P15 with a single dose of tamoxifen (resuspended to 20mg/ml in Corn Oil with 10% ethanol) by intraperitoneal injection, at a concentration of 0.15mg per gram of body weight. Pituitaries were harvested at the indicated time points post induction and processed for further analysis as described below. Mice were harvested from different litters for each time point at random. For litters in which there was a surplus of experimental mice, multiple samples were harvested for each required time point. For Wntless deletion studies, *Sox2*^*CreERT2/+*^;*Wls*^*fl/+*^;*ROSA26*^*mTmG/mTmG*^ males were bred with *Wls*^*fl/fl*^;*ROSA26*^*mTmG/mTmG*^ dams, to produce *Sox2*^*CreERT2/+*^;*Wls*^*fl/+*^;*ROSA26*^*mTmG/mTmG*^, *Sox2*^*CreERT2/+*^;*Wls*^*fl/fl*^;*ROSA26*^*mTmG/mTmG*^ and *Wls*^*fl/fl*^;*ROSA26*^*mTmG/mTmG*^ offspring. Pups of the indicated genotypes received intraperitoneal injections of 0.15mg of tamoxifen/gram body weight on 4 consecutive days, beginning at P14, and harvested 3 days after the final injection. For the β-catenin loss-of-function experiments, either *Sox2*^*CreERT2/+*^;*Ctnnb1*^*fl(ex2-6)/+*^;*ROSA26*^*mTmG/mTmG*^ or *Axin2*^*CreERT2/+*^;*Ctnnb1*^*fl(ex2-6)/+*^;*ROSA26*^*mTmG/mTmG*^ males were crossed with *Ctnnb1*^*fl(ex2-6)/ fl(ex2-6)/*^;*ROSA26*^*mTmG/mTmG*^ dams. *Axin2*^*CreERT2/+*^;*Ctnnb1*^*fl(ex2-6)/fl(ex2-6)*^;*ROSA26*^*mTmG/mTmG*^ and *Axin2*^*CreERT2/+*^;*Ctnnb1*^*fl(ex2-6)/+*^;*ROSA26*^*mTmG/mTmG*^ pups were induced with a single dose of tamoxifen, at a concentration of 0.15mg per gram of body weight and kept alive for 7 days before harvesting. *Sox2*^*CreERT2/+*^;*Ctnnb1*^*fl(ex2-6)/+*^;*ROSA26*^*mTmG/mTmG*^ and *Sox2*^*CreERT2/+*^;*Ctnnb1*^*fl(ex2-6)/fl(ex2-6)*^;*ROSA26*^*mTmG/mTmG*^ pups received two intraperitoneal injections of tamoxifen, at a concentration of 0.15mg/gram body weight, on two consecutive days and were kept alive for the indicated length of time before harvesting.

*TCF/LEF:H2B-EGFP/+* mice culled and the pituitaries harvested at the indicated ages for the respective experiments. For fluorescence-activated cell sorting experiments, mice were harvested at 21 days of age. *Axin2*^*CreERT2/+*^;*Sox2*^*eGFP/+*^*ROSA26*^*tdTomato/tdTomato*^ males were crossed with *ROSA26*^*tdTomat/tdTomato*^ dams to produce *Axin2*^*CreERT2/+*^;*Sox2*^*eGFP/+*^*ROSA26*^*tdTomato/tdTomato*^ that were induced with a single dose of tamoxifen at 21 days of age and harvested three days later for fluorescence-activated cell sorting experiments. For studies involving the induction of hypothyroidism, FVB/NJ mice were housed in a conventional facility on a 12 hour light/12 hour dark cycle and were given chow and water *ad libitum*. At weaning, P21 pups were either fed an iodine-deficient diet supplemented with 0.15% propylthiouracil (PTU) or a normal maintenance diet for control animals. Following 7 days of treatment, pituitaries were collected and fixed in 10% NBF for 18 hours at room temperature.

### Flow cytometry analysis of lineage traced pituitaries

For the quantification of cells by flow cytometry, anterior lobes of *Axin2*^*CreERT2/+*^;*ROSA26*^*mTmG/+*^ mice dissected at the indicated time points. The posterior and intermediate lobes were dissected from the anterior lobes under a dissection microscope. Untreated *ROSA26*^*mTmG/+*^ and wild type pituitaries from age-matched litters were used as tdTomato only and negative controls, respectively. Dissected pituitaries were incubated in Enzyme Mix (0.5% w/v collagenase type 2 (Lorne Laboratories), 0.1x Trypsin (Gibco), 50αg/ml DNase I (Worthington) and 2.5αg/ml Fungizone (Gibco) in Hank’s Balanced Salt Solution (HBSS)(Gibco)) in a cell culture incubator for up to 3 hours. 850ml of HBSS were added to each Eppendorf in order to quench the reaction. Pituitaries were dissociated by agitation, pipetting up and down 100x at first with a 1ml pipette, followed by 100x with a 200αl pipette. Cells were transferred to a 15ml Falcon tube and resuspended in 9ml of HBSS and spun down at 200g for 5 minutes. The supernatant was aspirated, leaving behind the cell pellet that was resuspended in PBS and spun down at 1000rpm for 5 minutes before being resuspended in a Live/Dead dye (Life Technologies, L34975) prepared to manufacturer’s instructions, for 30 minutes in the dark. Cells were washed in PBS as above. The pellet was resuspended in FIX & PERM Cell Permeabilization Kit (Life Technologies, GAS003) prepared as per manufacturer’s instructions for 10 minutes at room temperature. Cells were washed as above, and the pellet was resuspended in 500µl of FACS buffer (1% fetal calf serum (Sigma), 25mM HEPES in PBS) and filtered through 70αm filters (BD Falcon), into 5ml round bottom polypropylene tubes (BD Falcon). 1 minute prior to analysis, 1µl of Hoechst was added to the suspended cells and incubated. Samples were analysed on a BD Fortessa, and gated according to negative and single fluorophore controls. Single cells were gated according to SSC-A and SSC-W. Dead cells were excluded according to DAPI (2ng/ml, incubated for 2 mins prior to sorting). GFP^+^, tdTomato^+^ and GFP^+^;tdTomato^+^ cells were gated according to negative controls in the PE-A and FITC-A channels.

### Fluorescence Activated Cell Sorting for sequencing or colony forming assays

For fluorescence activated cell sorting, the anterior lobes from *Sox2*^*eGFP/+*^, TCF/LEF:H2B-GFP or *Axin2*^*CreERT2/+*^;*ROSA26*^*tdTomato*^;*Sox2*^*eGFP/+*^ and their respective controls were dissected and dissociated as above. After dissociation cells were spun down at 200g in HBSS and the pellet was resuspended in 500µl FACS buffer. Using an Aria III FACs machine (BD systems), samples were gated according to negative controls, and where applicable single fluorophore controls. Experimental samples were sorted according to their fluorescence, as indicated, into tubes containing either RNAlater (Qiagen) for RNA isolation or 1ml of Pit Complete Media for culture ((Pit Complete: 20ng/ml bFGF and 50ng/ml of cholera toxin in ‘Pit Basic’ media (DMEM-F12 with 5% Fetal Calf Serum,100U/ml Penicillin and 100αg/ml Streptomycin). Cells were plated in 12-well plates at clonal density, approximately 500 cells/well. Colonies were incubated for 7 days total before being fixed in 10% neutral buffered formalin (NBF) (Sigma) for 10 minutes at room temperature, washed for five minutes, three times, mins with PBS and stained with crystal violet in order for the number of colonies to be quantified.

### RNA-sequencing

Total RNA was isolated from each sample and following poly-A selection, cDNA libraries were generated using TruSeq (Clontech, 634925). Barcoded libraries were then pooled at equal molar concentrations and sequenced on an Illumina Hiseq 4000 instrument in a 75 base pair, paired – end sequencing mode, at the Wellcome Trust Centre for Human Genetics (Oxford, United Kingdom). Raw sequencing reads were quality checked for nucleotide calling accuracy and trimmed accordingly to remove potential sequencing primer contaminants. Following QC, forward and reverse reads were mapped to GRCm38/mm10 using Hisat2 (Kim et al., 2015). Using a mouse transcriptome specific GTF as a guide, FeatureCounts (Liao et al., 2014) was used to generate gene count tables for every sample. These were utilised within the framework of the Deseq2 (Love et al., 2014) and FPKM values (generated by FPKM count (Wang et al., 2012)) were processed using the Cufflinks (Trapnell et al., 2012) pipelines which identified statistically significant gene expression differences between the sample groups. Following identification of differentially expressed genes (at an FDR < 0.05) we focused on identifying differentially expressed pathways using a significance threshold of FDR < 0.05 unless otherwise specified. The gene lists used for Gene Set Enrichment Analysis (GSEA) were as found on the BROAD institute GSEA MSigDBv.7 ‘molecular signatures database’. The deposited dataset can be accessed through the following link: https://dataview.ncbi.nlm.nih.gov/object/PRJNA421806?reviewer=kr90aklsdtikh3gkh3tdlpv30s

### Immunofluorescence and microscopy

Freshly harvested pituitaries were washed in PBS for 10 minutes before being fixed in 10% NBF for 18 hours at room temperature. In short, embryos and whole pituitaries were washed in PBS 3 times, before being dehydrated through a series of 1 hour washes in 25%, 50%, 70%, 80%, 90%, 95% and 100% ethanol. Tissues were washed in Neo-Clear (Sigma) at room temperature for 10 minutes, then in fresh preheated Neo-Clear at 60 **°**C for 10 minutes. Subsequently, a mixture of 50% Neo-Clear:50% paraffin wax at 60**°**C for 15 minutes followed by three changes of pure wax for a minimum of 1 hour washes at 60**°**C, before being orientated to be sectioned in the frontal plane. Embedded samples were sectioned at 5µm and mounted on to Super Frost+ slides.

For immunofluorescence, slides were deparaffinised in Neo-Clear for three times ten minutes, washed in 100% ethanol for three times five minutes, and rehydrated in a series of five minute ethanol washes up to distilled water (95%, 90%, 80%, 70%, 50%, 25%, H2O). Heat induced epitope retrieval was performed with 1x DeClear Buffer (citrate pH 6) in a Decloaking chamber NXGEN (Menarini Diagnostics) for 3 minutes at 110°C. Slides were left to cool to room temperature before proceeding to block for 1 hour at room temperature in Blocking Buffer (0.2% BSA, 0.15% glycine, 0.1% TritonX in PBS) with 10% serum (sheep or donkey, depending on secondary antibodies). Primary antibodies were diluted in blocking buffer with 1% of the appropriate serum and incubated overnight at 4**°**C. Slides were washed three times for 10 minutes with PBST. Slides were incubated with secondary antibodies diluted 1:400 in blocking buffer with 1% serum for one hour at room temperature. Slides were washed three times with PBST as above. Where biotinylated secondary antibodies were used, slides were incubated with streptavidin diluted 1:400 in blocking buffer with 1% serum for one hour at room temperature. Finally, slides were washed with PBST and mounted using Vectashield Antifade Mounting Medium (Vector Laboratories, H-1000).

The following antibodies, along with their dilutions and detection technique, were used: GFP (1:400, Alexa Fluor-488 or −647 secondary), SOX2 raised in goat (1:200, Alexa Fluor-488 secondary), SOX2 raised in rabbit (1:100, biotinylated secondary), SOX9 (1:500, biotinylated secondary), PIT1 (1:500, biotinylated secondary), SF1 (1:300, biotinylated secondary), TPIT (1:200, biotinylated secondary), Ki-67 (1:100, biotinylated secondary), pH-H3 (1:500, biotinylated secondary), GH (1:1000, biotinylated secondary), TSH (1:1000, biotinylated secondary), PRL (1:1000, biotinylated secondary), ACTH (1:400, Alexa Fluor-555 secondary), LH/FSH (1:300, biotinylated secondary), ZO-1 (1:300, Alexa Fuor-488), E-Cadherin (1:300, Alexa Fluor-488). Nuclei were visualized with Hoechst (1:1000). Images were taken on a TCS SPS Confocal (Leica Microsystem) with a 20x objective for analysis.

### mRNA *In Situ* Hybridisation

All mRNA *in situ* hybridisations were performed using the RNAscope singleplex or duplex chromogenic kits (Advanced Cell Diagnostics) on formalin fixed paraffin embedded sections processed as described in the above section. The protocol followed the manufacturer’s instructions with slight modifications. ImmEdge Hydrophobic Barrier PAP Pen (Vector Laboratories, H-4000) was used to draw a barrier around section while air-drying following the first ethanol washes. Pretreatment followed the standard length of time for pituitaries (twelve minutes), while embryos were boiled for 10 minutes. For singleplex, the protocol proceeded to follow the instructions exactly. For duplex, Amplification 9 was extended to one hour and the dilution of the Green Detection reagent was increased to 1:30. For both protocols, sections were counterstained with Mayer’s Haematoxylin (Vector Laboratories, H-3404), left to dry at 60**°**C for 30 minutes before mounting with VectaMount Permanent Mounting Medium (Vector Laboratories, H-5000). Slides were scanned using a Nanozoomer-XR Digital Slide Scanner (Hamamatsu) and processed using Nanozoomer Digital Pathology View (Hamamatsu).

### Quantification of cells

Cell numbers were quantified in ImageJ using the cell counter plugin (Schindelin et al., 2012). At a minimum, three sections per pituitary were quantified, spaced no less than 100µM apart in the tissue.

### Statistics

All statistical analyses were performed in GraphPad Prism. Data points in graphs represent the mean values of recordings from a single biological replicate unless otherwise stated.

## ACKNOWLEDGEMENTS

This study has been supported by the Medical Research Council (MR/L016729/1, MR/T012153/1) (C.L.A.), The Lister Institute of Preventive Medicine (C.L.A.), the Deutsche Forschungsgemeinschaft (DFG German Research Foundation) (Project Number 314061271 – TRR 205) (C.L.A.), the Howard Hughes Medical Institute (R.N.), the Agence Nationale de la Recherche (ANR-18-CE14-0017) and Fondation pour la Recherche Médicale (DEQ20150331732) (P.M.). J.P.R. was supported by a Dianna Trebble Endowment Fund Dental Institute Studentship, E.J.L. by the King’s Bioscience Institute and the Guy’s and St Thomas’ Charity Prize PhD Programme in Biomedical and Translational Science, Y.K. by a Project Support Grant from the British Society for Neuroendocrinology. We thank Dr A.F. Parlow and the National Hormone and Peptide Program (Harbor–University of California, Los Angeles Medical Center) for providing some of the antibodies used in this study and Prof. J. Drouin and Prof. S. Rhodes for TPIT and PIT1 antibodies respectively. We thank the High-Throughput Genomics Group at the Wellcome Trust Centre for Human Genetics (funded by Wellcome Trust grant reference 090532/Z/09/Z) for the generation of the Sequencing data. For flow sorting and analysis, this research was supported by the National Institute for Health Research (NIHR) Biomedical Research Centre based at Guy’s and St Thomas’ NHS Foundation Trust and King’s College London. We thank Marie Isabelle Garcia, Juan Pedro Martinez-Barbera and Paul Le Tissier for useful discussions and critical comments on the manuscript.

## AUTHOR CONTRIBUTIONS

Conceptualization C.L.A. and J.P.R.; Methodology J.P.R., C.L.A., P.M.; Investigation J.P.R., V.Y., A.S., E.J.L., Sh.Ha., Sc.Ha., C.L.A., Y.K.; Resources R.N., B.W., M.F., X.L., Y.K., P.M.; Writing – Original Draft, C.L.A. and J.P.R.; Writing-Review & Editing C.L.A., J.P.R. R.N., X.L., P.M.; Supervision C.L.A., R.N.; Funding Acquisition C.L.A., R.N., P.M.

## KEY RESOURCES TABLE

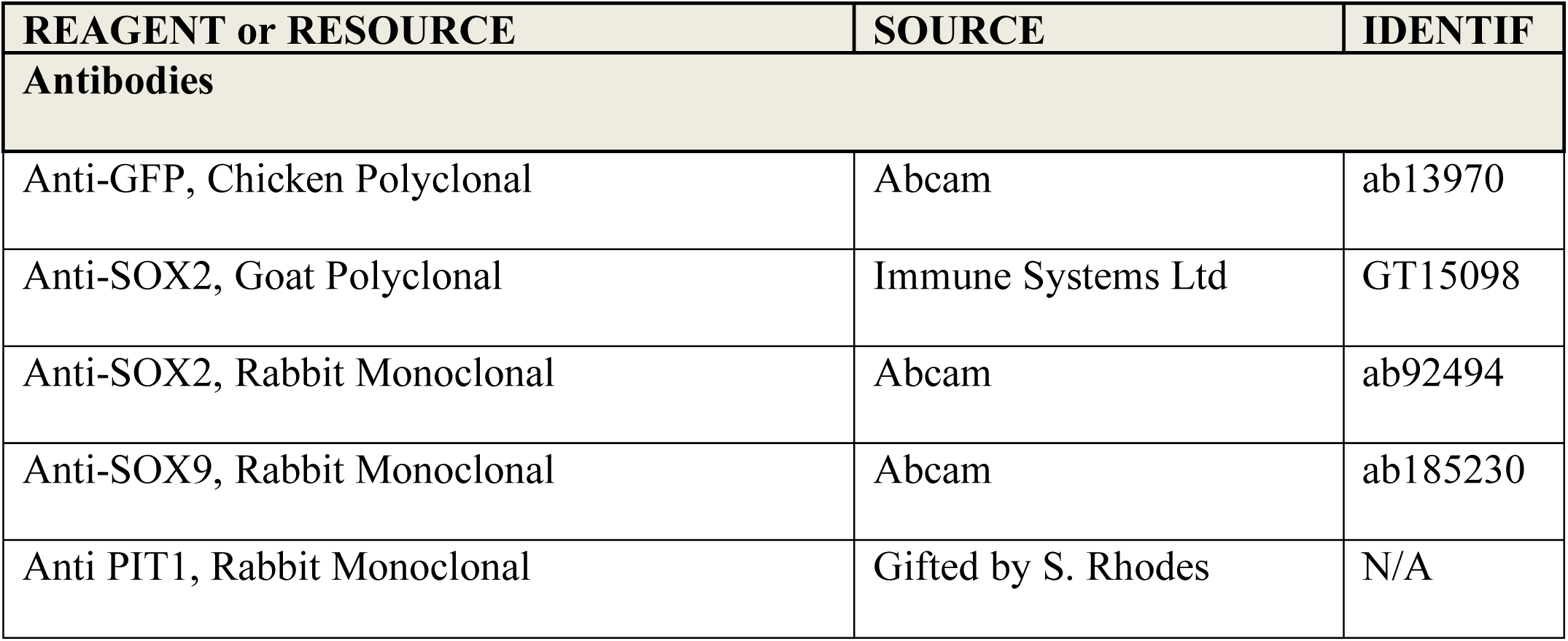
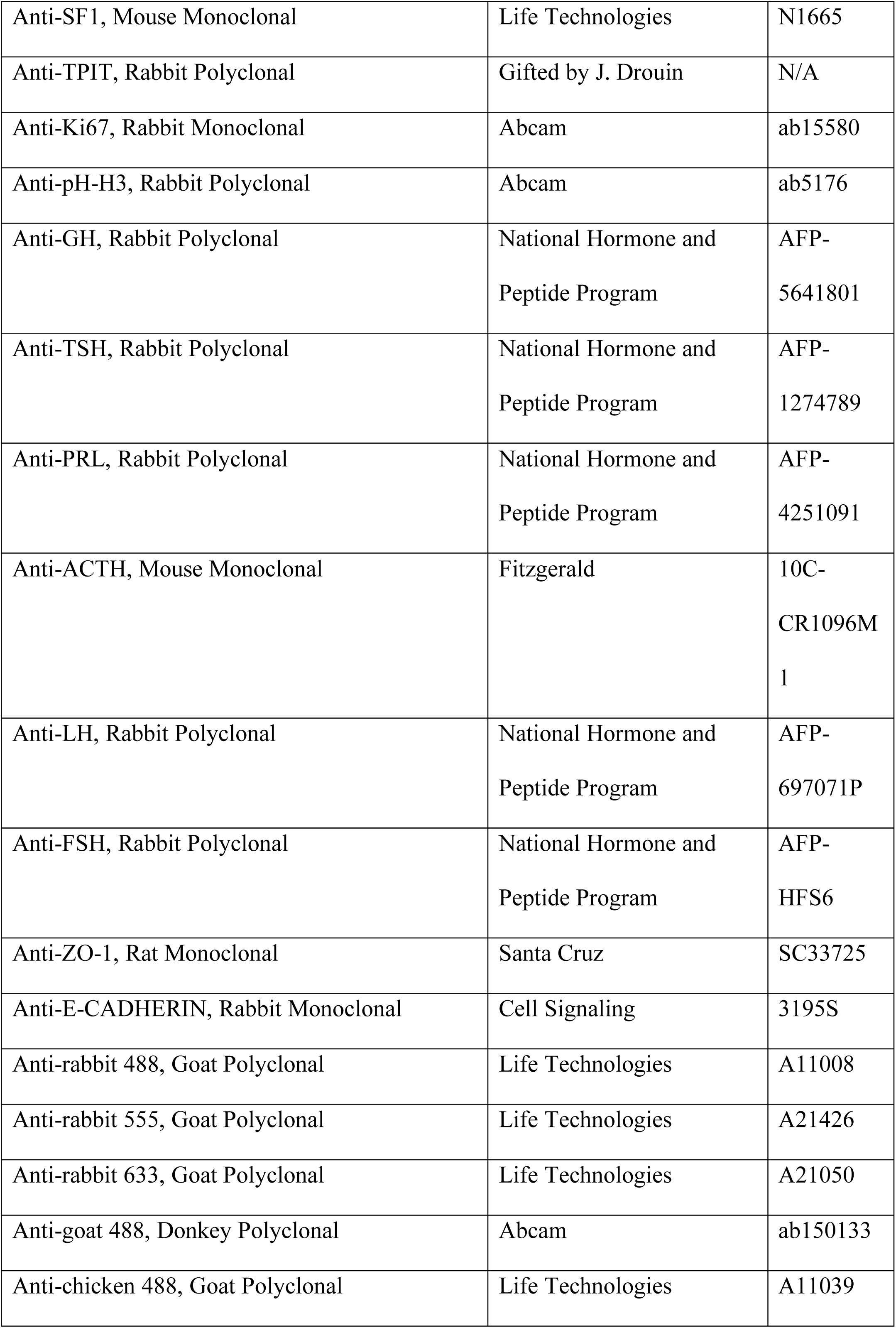
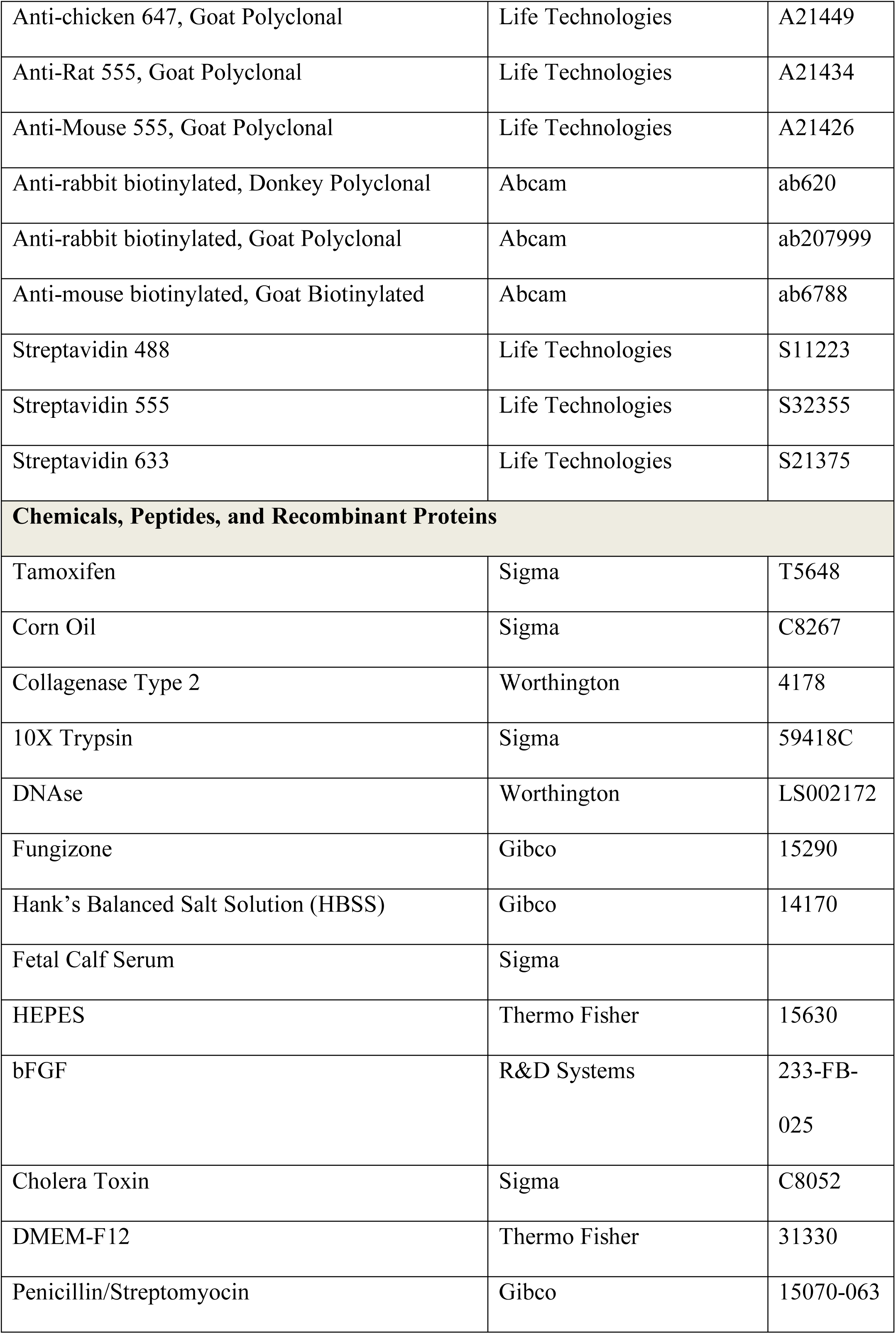
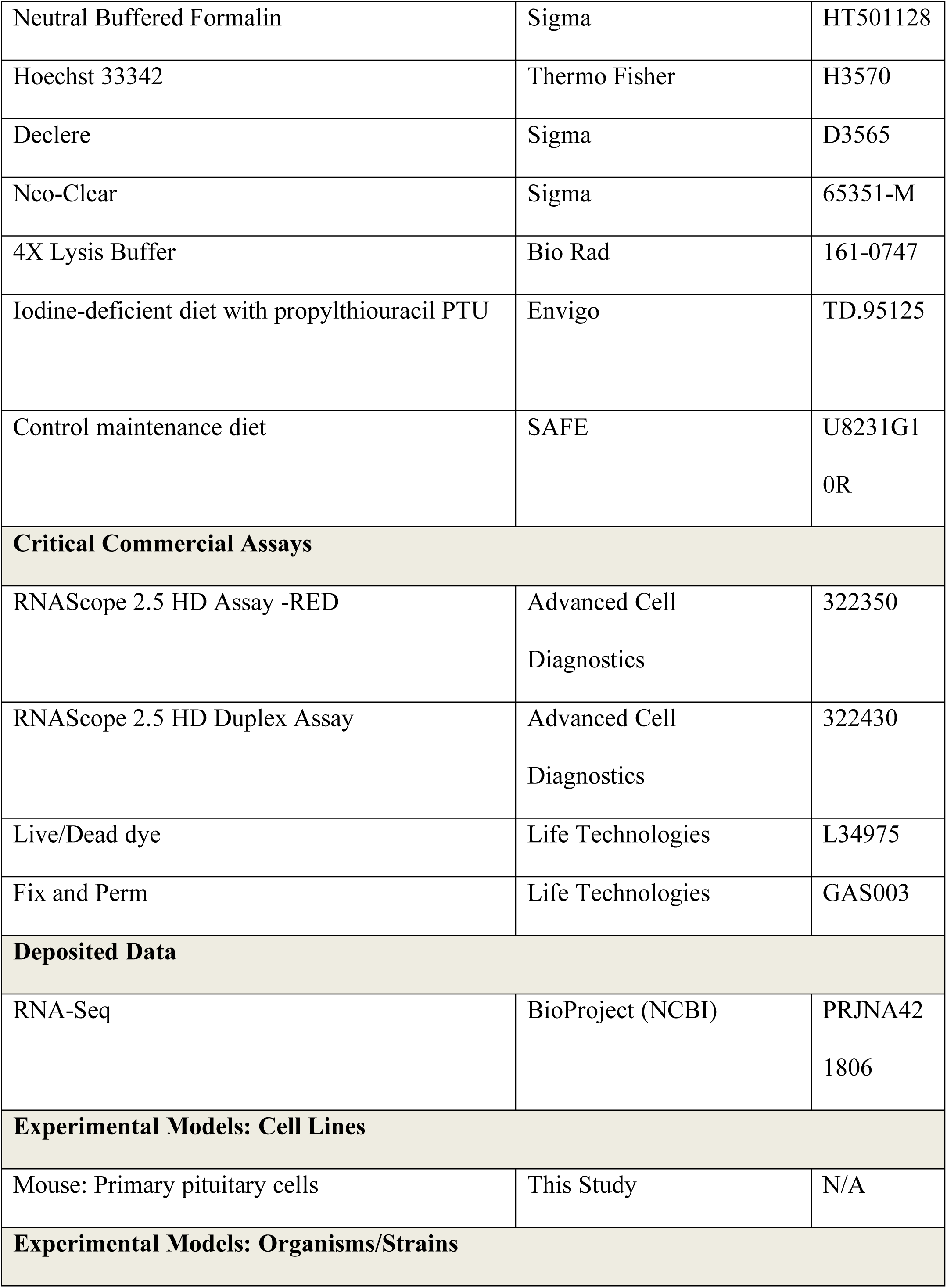
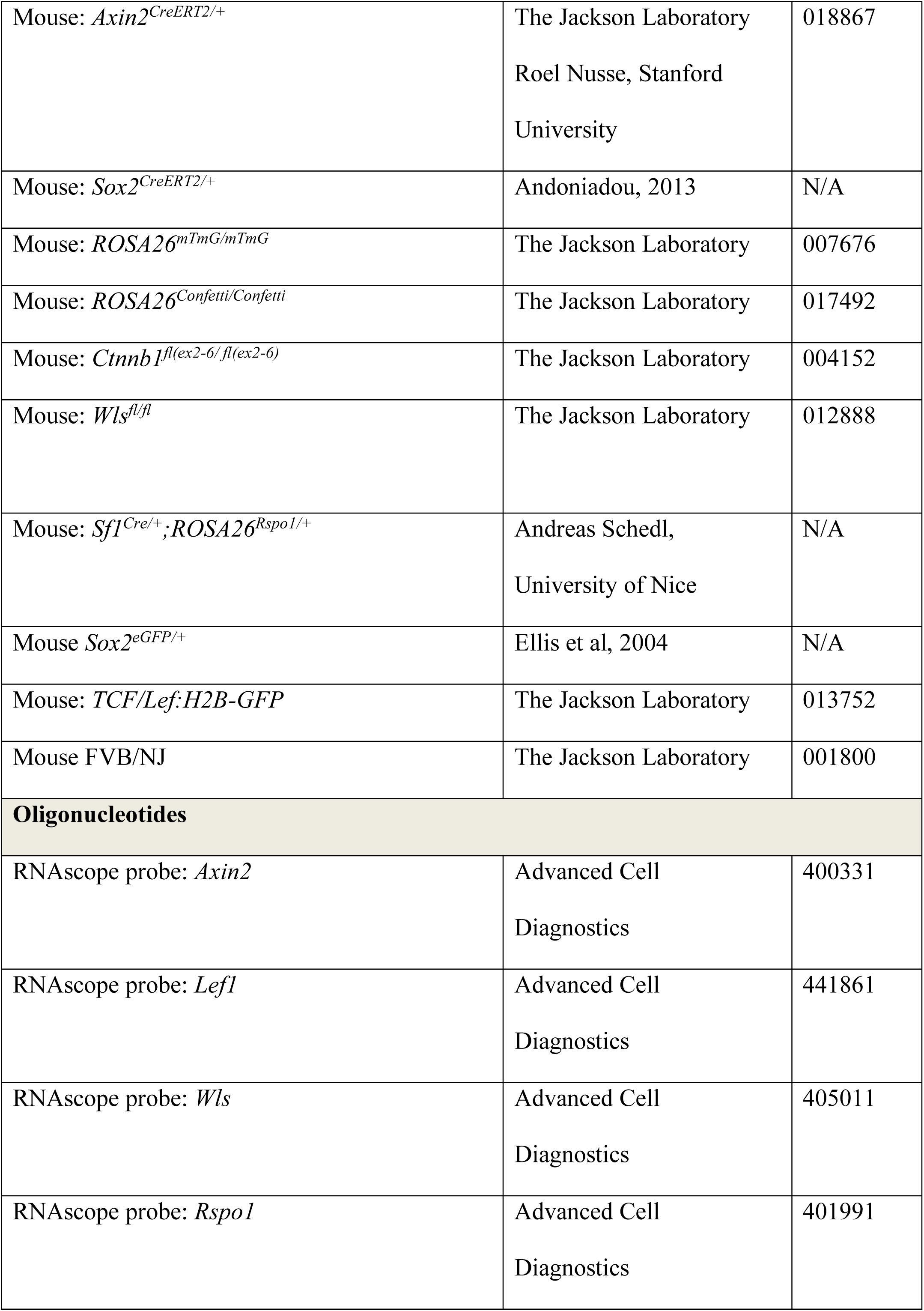
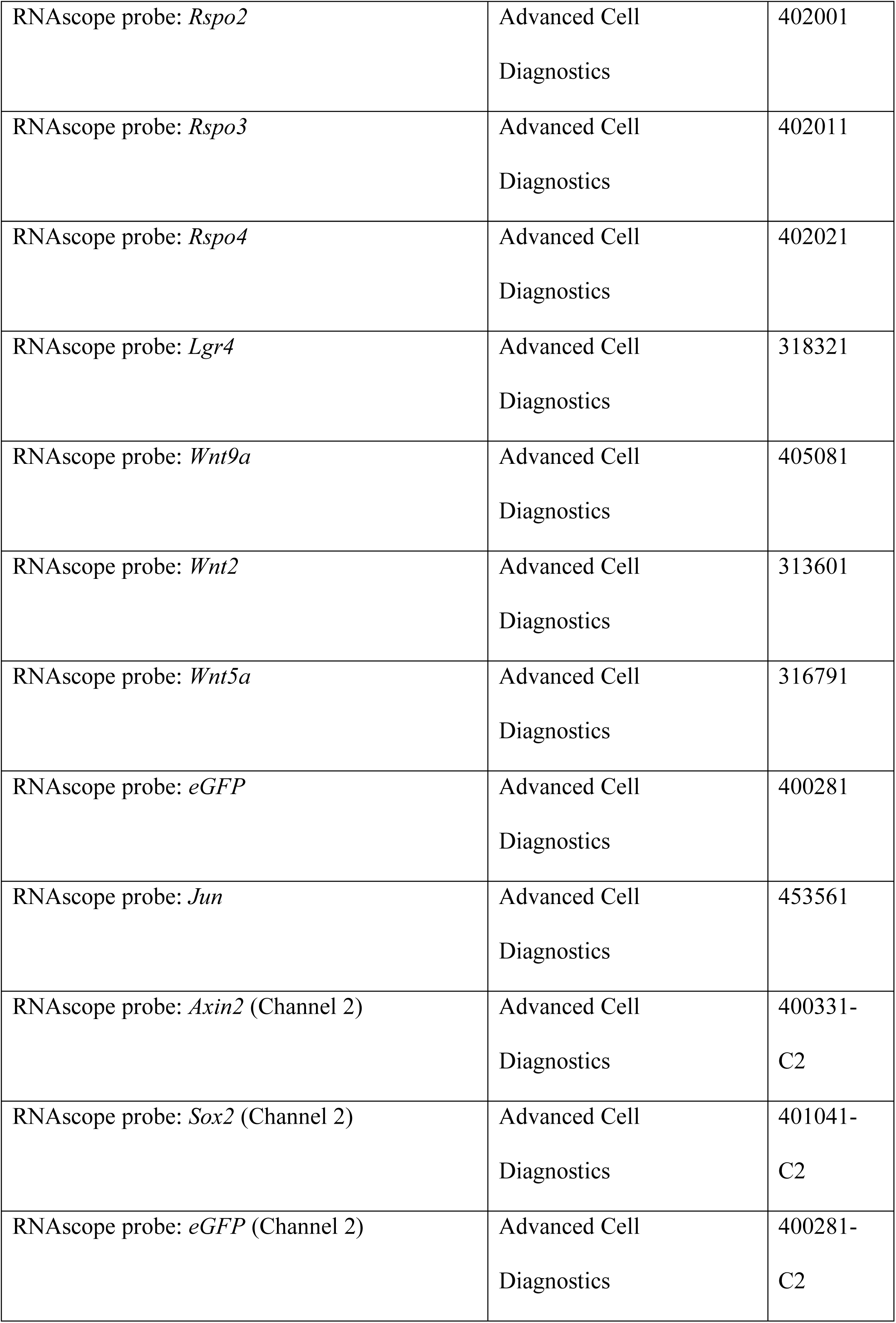
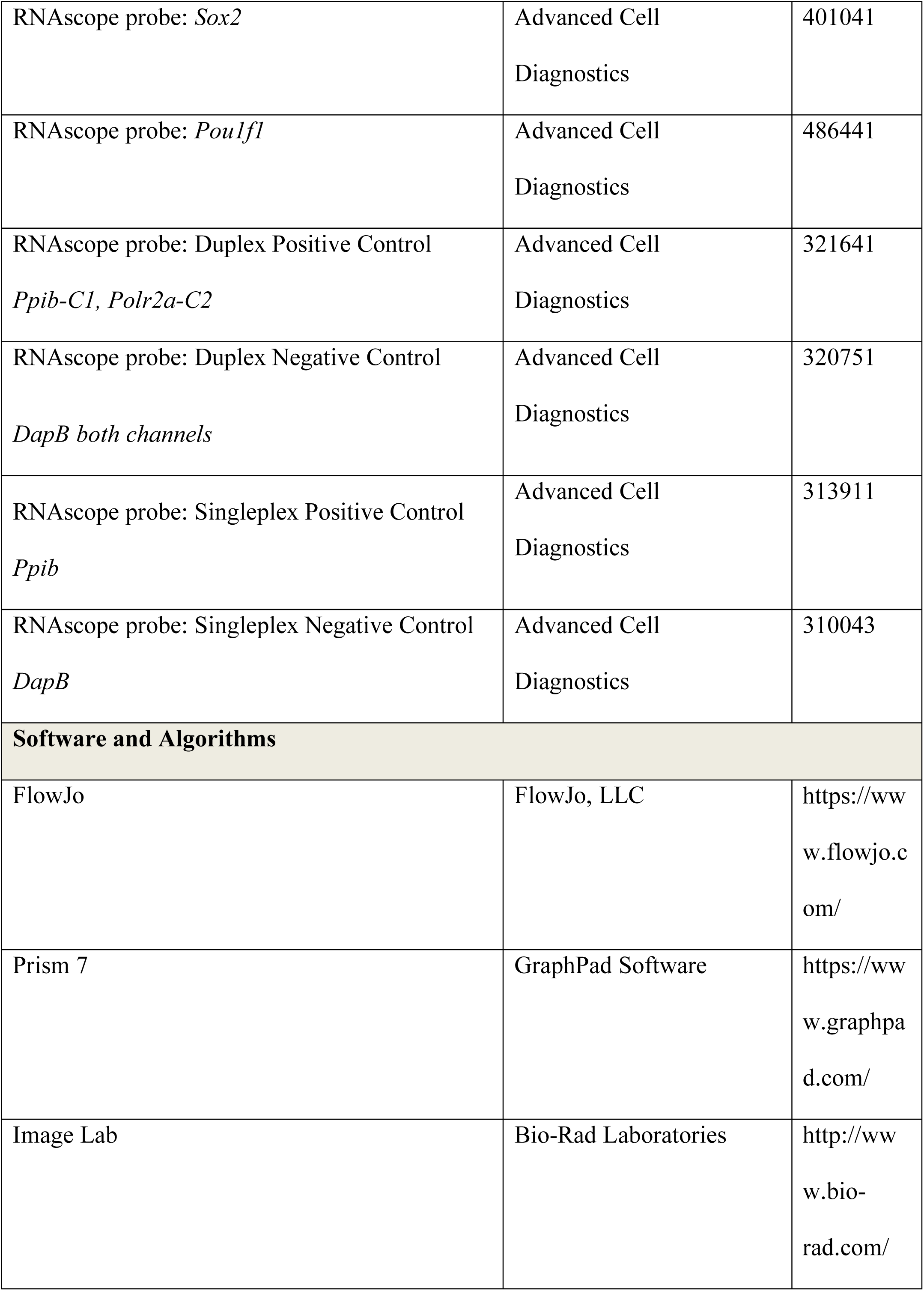
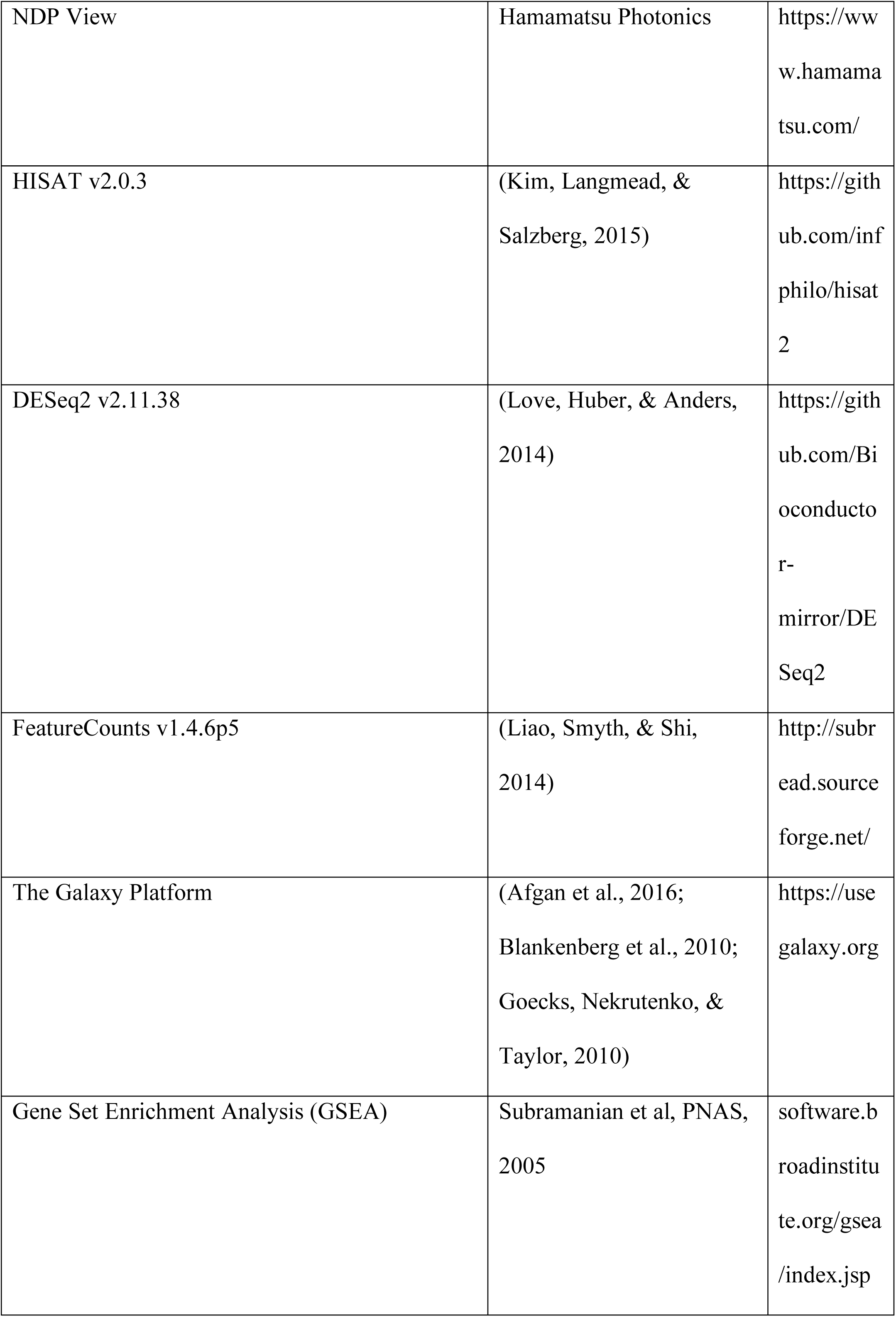
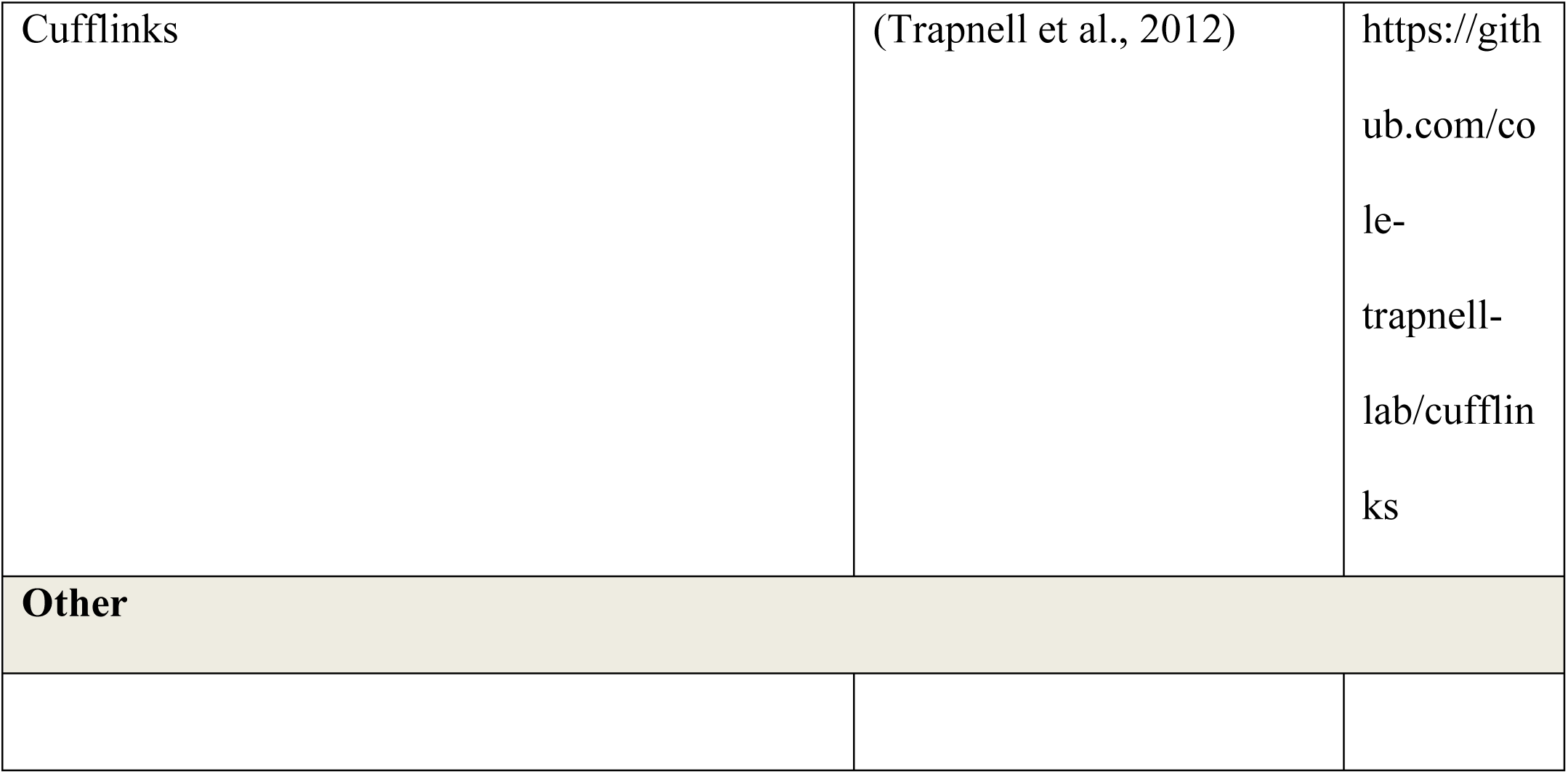

