## Supplementary Figure for "Pituitary stem cells produce paracrine WNT signals to control the expansion of their descendant progenitor cells"

### SUPPLEMENTARY INFORMATION

### TABLES

**Supplementary Table 1. Gene lists of Gene Set Enrichment Analyses**

### SUPPLEMENARY FIGURES

**Supplementary Figure 1. *Axin2* expressing cells contribute to pituitary growth and expansion of all lineages**

**A.** Immunofluorescence staining against GFP (green) and markers of hormone-secreting endocrine cells (GH (somatotrophs), ACTH (corticotrophs), PRL (lactotrophs), TSH (thyrotrophs), FSH/LH (gonadotrophs)) in $Axin2^{CreERT2/+};Rosa26^{mTmG/+}$ pituitaries induced at P14 and lineage traced for 48 hours. Scale bar 10µm.

B. Clonal analysis of individual cells targeted in $Sox2^{CreERT2/+};Rosa26^{Confetti/+}$ (left panel) and $Axin2^{CreERT2/+};Rosa26^{Confetti/+}$ pituitaries (right panel), induced at P14 and harvested after 4 weeks (P42). Arrows point to individual clones, numbered for the number of cells in the clone. Scale bar 100µm.

C. Immunofluorescence staining against TSH (red) in pituitaries from mice that were fed with a normal diet or with low iodine diet supplemented with 0.15% PTU for 7 days, inducing hypothyroidism. Quantification of the number of $TSH^+$ cells per section. Values represent mean +/- SEM, $P<0.0001$, unpaired $t$-test with Welch's correction ($n = 3$ pituitaries). Scale bar 50µm.

D. Graph of the percentage of pH-H3 positive cells in pituitaries of mice fed on normal or low iodine diet, showing a significant increase in diving cells (in G2/M) in hypothyroid animals. Values represent mean +/- SEM, $P=0.0308$, unpaired $t$-test ($n = 3$ pituitaries per condition).

E. mRNA *in situ* hybridisation using specific probes against *Axin2* (red) in pituitaries of mice fed on normal or low iodine diet, detecting an increase in transcripts in hypothyroid animals. Brightfield and native FastRed fluorescence images shown. Quantification of *Axin2* fluorescent signal intensity. Values represent mean +/- SEM, *P*= 0.0378, upaired *t*-test with Welch's correction (*n* = 3 pituitaries per condition).

F. Dorsal wholemount view of $Axin2^{CreERT2/+}$; $Ctnnb1^{LOF/+}$;$Rosa26^{mTmG/+}$and $Axin2^{CreERT2/+}$; $Ctnnb1^{LOF/LOF}$;$Rosa26^{mTmG/+}$ pituitaries induced at P14 and lineage traced for 5 days. Immunofluorescence staining against GFP (green) and pH-H3 (magenta) in $Axin2^{CreERT2/+}$; $Ctnnb1^{LOF/+}$;$Rosa26^{mTmG/+}$and $Axin2^{CreERT2/+}$; $Ctnnb1^{LOF/LOF}$;$Rosa26^{mTmG/+}$ pituitaries. Scale bar 50µm. Quantification of the contribution of lineage traced cells in control and mutants. Each data point represents the mean from one individual. *P*=0.0313, unpaired *t*-test (n = 3).

Supplementary Figure 1

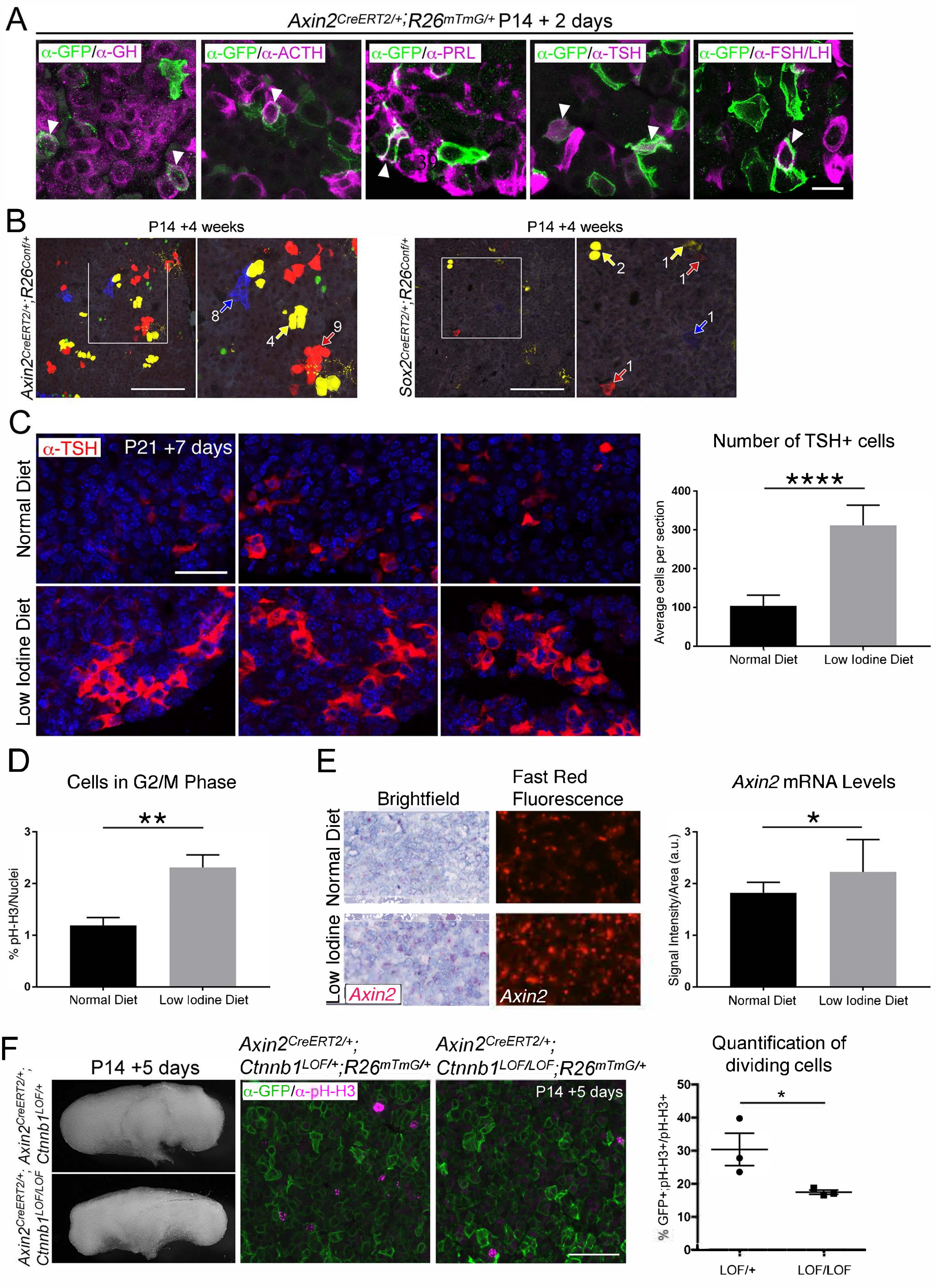

**Supplementary Figure 2. Activation of WNT signalling in SOX2$^{+}$ PSCs and their descendants is necessary for long-term growth**

A. Confocal images of native GFP fluorescence in frontal sections from *TCF/Lef:H2B-EGFP* pituitaries at P21. Scale bar 50µm.

B. mRNA *in situ* hybridisation in *TCF/Lef:H2B-EGFP* pituitaries at P21, detecting *Egfp* transcripts (red). Double mRNA *in situ* hybridisation showing overlap between *Sox2* (red) and *Egfp* (blue) transcripts in pituitaries at P21. White arrowheads indicate double-positive staining. Scale bars 50µm.

C. Immunofluorescence staining against SOX2 (magenta) and GFP (green) in *TCF/Lef:H2B-EGFP* pituitaries harvested from P21 mice. White arrows indicate double positive cells. Representative scatter plot showing gating used for fluorescence activated cell sorting and population percentages in each gate. Graph of quantification of the *in vitro* colony forming potential of GFP cells isolated from P21 *TCF/Lef:H2B-EGFP* pituitaries by flow sorting. Each data point represents single well replicates. Error bars show SEM, $P<0.001$ (One-way ANOVA, n = 3 experiments). Scale bar 50µm.

D. Immunofluorescence staining against PIT1, TPIT and SF1 (magenta) in $Sox2^{CreERT2/+}$;$Ctnnb1^{LOF/+}$;$Rosa26^{mTmG/+}$ and $Sox2^{CreERT2/+}$;$Ctnnb1^{LOF/LOF}$;$Rosa26^{mTmG/+}$ pituitaries 22 weeks post-induction at P14 (age P24). Arrows indicate double positive cells. Scale bar 50µm.

E. Immunofluorescence staining against β-catenin (magenta) and GFP (green) in $Sox2^{CreERT2/+}$; $Ctnnb1^{LOF/+}$;$Rosa26^{mTmG/+}$ and $Sox2^{CreERT2/+}$; $Ctnnb1^{LOF/LOF}$;$Rosa26^{mTmG/+}$ pituitaries 22 weeks post-induction. Arrowheads indicate double positive cells, arrows indicate GFP$^{+}$ cells that have lost β-catenin expression in mutants. Scale bar 50µm.

PL, posterior lobe; IL, Intermediate lobe; AL, anterior lobe; Inf, infundibulum; RP, Rathke's pouch; Sph, sphenoid bone.

Supplementary Figure 2

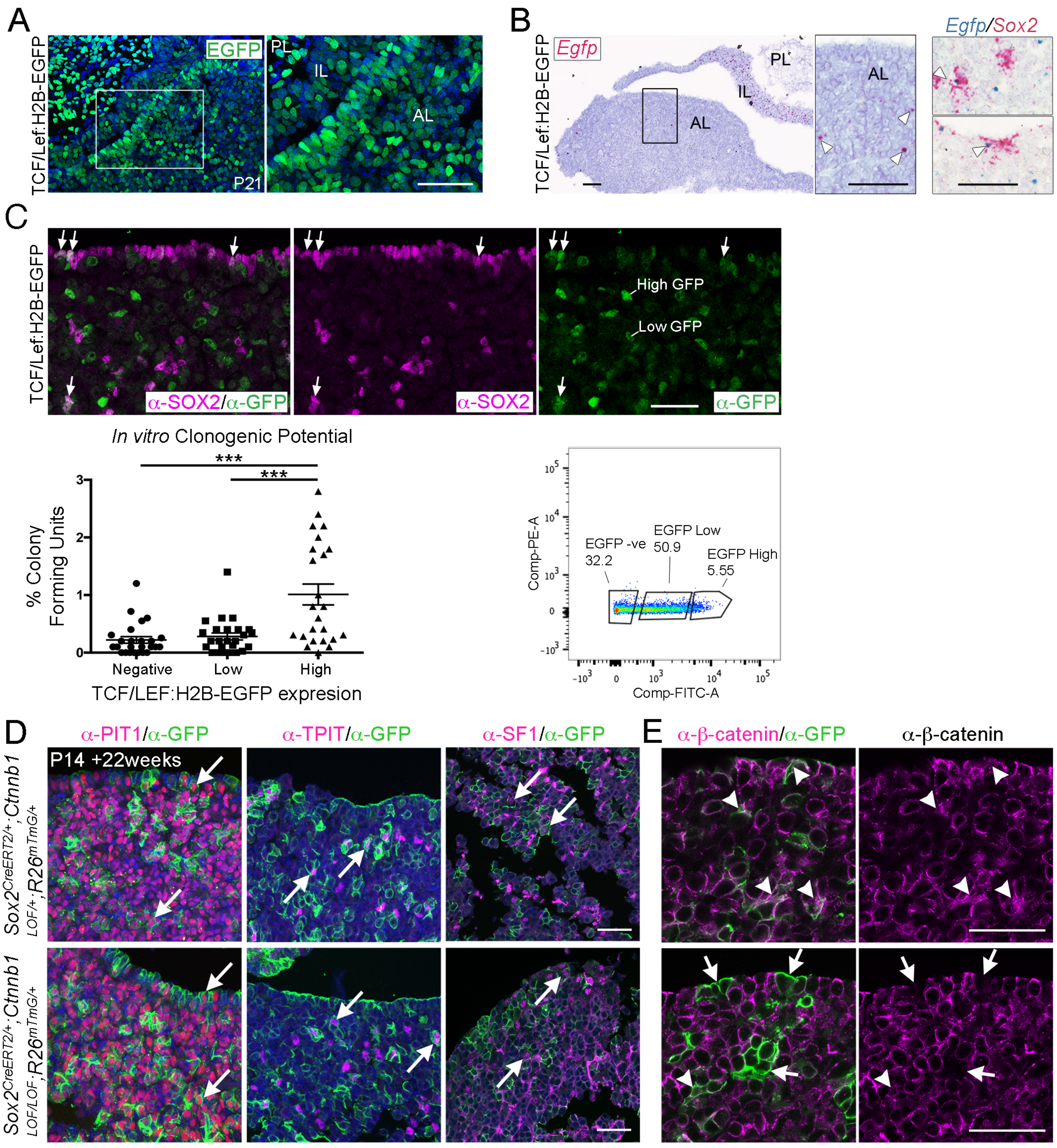

**Supplementary Figure 3. SOX2$^{+}$ PSCs are as a source of WNT ligands in the pituitary**

A. Native EGFP protein expression in frontal cryosection of a P14 *Sox2*$^{Egfp/+}$ pituitary. Schematic of the workflow used for bulk RNA-sequencing analysis of *Sox2*$^{+}$ and *Sox2*$^{-}$ cells. Genome browser views of reads aligning to the *Sox2* and *Pit1* loci in the positive and negative fractions indicating good separation of the EGFP$^{+}$ population. Scale bar 50μm.

B. *Sox2*$^{+}$ cells express a significant enrichment in markers associated with epithelial-to-mesenchymal transition (EMT), adherens and tight junctions, consistent with their epithelial nature. GSEA plots and immunofluorescence staining against E-Cadherin (adherens junction marker) and ZO1 (tight junction marker) in the marginal zone epithelium at P14. Scale bar 50μm. See Supplementary Table 1 for full GSEA gene lists.

C. *Sox2*$^{+}$ cells express a significant enrichment in several signalling pathways, shown with respective GSEA plots.

D. Bar charts showing the FPKM values of components of the LGR/RNF43/ZNRF3/RSPONDIN module in the *Sox2+* and *Sox2-* fractions and the distribution of the Frizzled receptors. GSEA plot for components of the WNT pathway. Validation of sequencing: (i) mRNA *in situ* hybridisation with specific probes against *Lgr4* (blue) and *Sox2* (red) in P14 pituitaries showing co-expression. (ii) Double mRNA in situ hybridisation against *Fzd4* (blue) and *Sox2* (red) indicating co-expression in both the marginal zone epithelium and parenchymal *Sox2*$^{+}$ cells. Boxed regions are magnified. Scale bars 250μm and 50μm in boxed inserts. (iii) mRNA in situ hybridisation against *Rspo1*, *Rspo2*, *Rspo3* and *Rspo4* in sagittal sections of wild type pituitaries at P14. Boxed regions are magnified, only *Rspo4* is detected. Scale bars 250µm and 100µm in boxed inserts.

Supplementary Figure 3

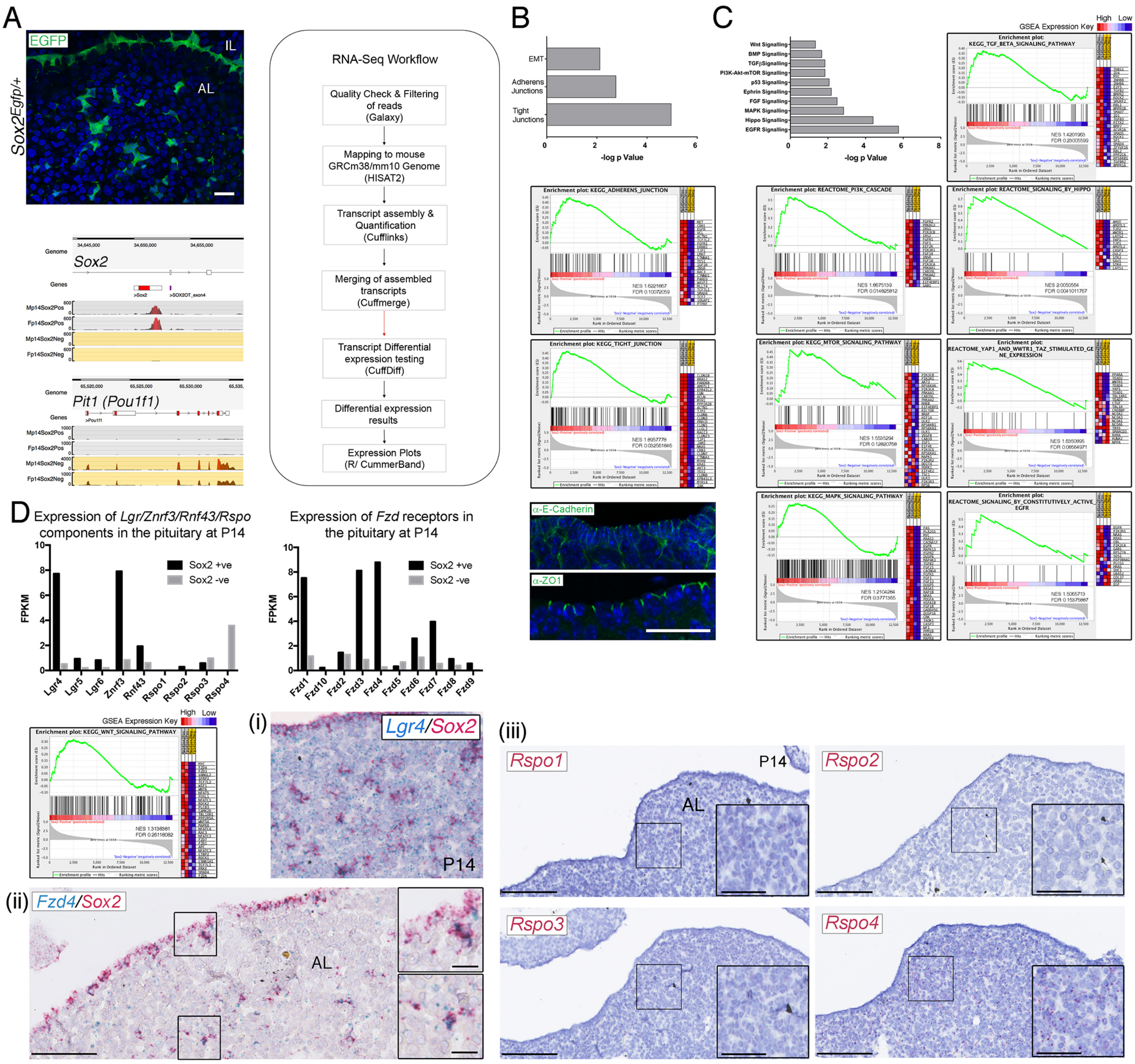

**Supplementary Figure 4. Paracrine secretion of WNTs from SOX2$^{+}$ PSCs is necessary for expansion of committed cells**

A. Schematic of time points for induction by tamoxifen induction and tissue harvesting of control *Sox2$^{+/+}$;Wls$^{fl/fl}$* and mutant *Sox2$^{CreERT2/+}$;Wls$^{fl/fl}$* pituitaries.

B. Whole mount, dorsal views of control *Sox2$^{+/+}$;Wls$^{fl/fl}$* (top panel) and mutant *Sox2$^{CreERT2/+}$;Wls$^{fl/fl}$* (bottom panel) pituitaries at P21.

Supplementary Figure 4

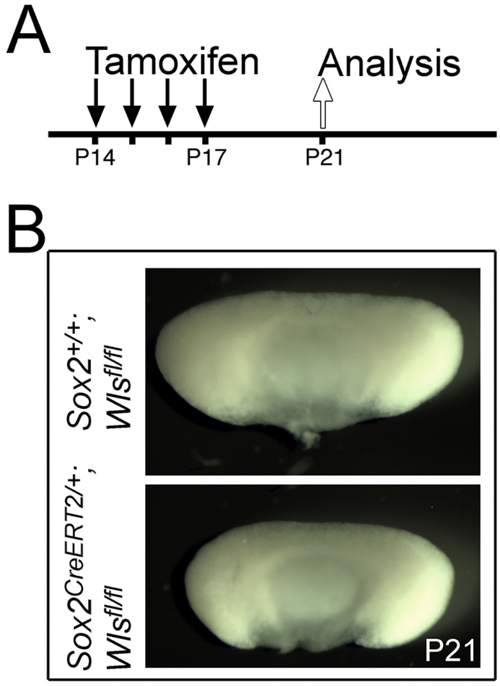
